# Population Dynamics of EMT Elucidates the Timing and Distribution of Phenotypic Intra-tumoral Heterogeneity

**DOI:** 10.1101/2023.01.13.523978

**Authors:** Annice Najafi, Mohit K. Jolly, Jason T. George

## Abstract

The Epithelial-to-Mesenchymal Transition (EMT) is a hallmark of cancer metastasis and morbidity. EMT is a non-binary process, and cells can be stably arrested en route to EMT in an intermediate hybrid state associated with enhanced tumor aggressiveness and worse patient outcomes. Understanding EMT progression in detail will provide fundamental insights into the mechanisms underlying metastasis. Despite increasingly available single-cell RNA sequencing data that enable in-depth analyses of EMT at the single-cell resolution, current inferential approaches are limited to bulk microarray data. There is thus a great need for computational frameworks to systematically infer and predict the timing and distribution of EMT-related states at single-cell resolution. Here, we develop a computational framework for reliable inference and prediction of EMT-related trajectories from single-cell RNA sequencing data. Our model can be utilized across a variety of applications to predict the timing and distribution of EMT from single-cell sequencing data.

**Graphical Abstract:** 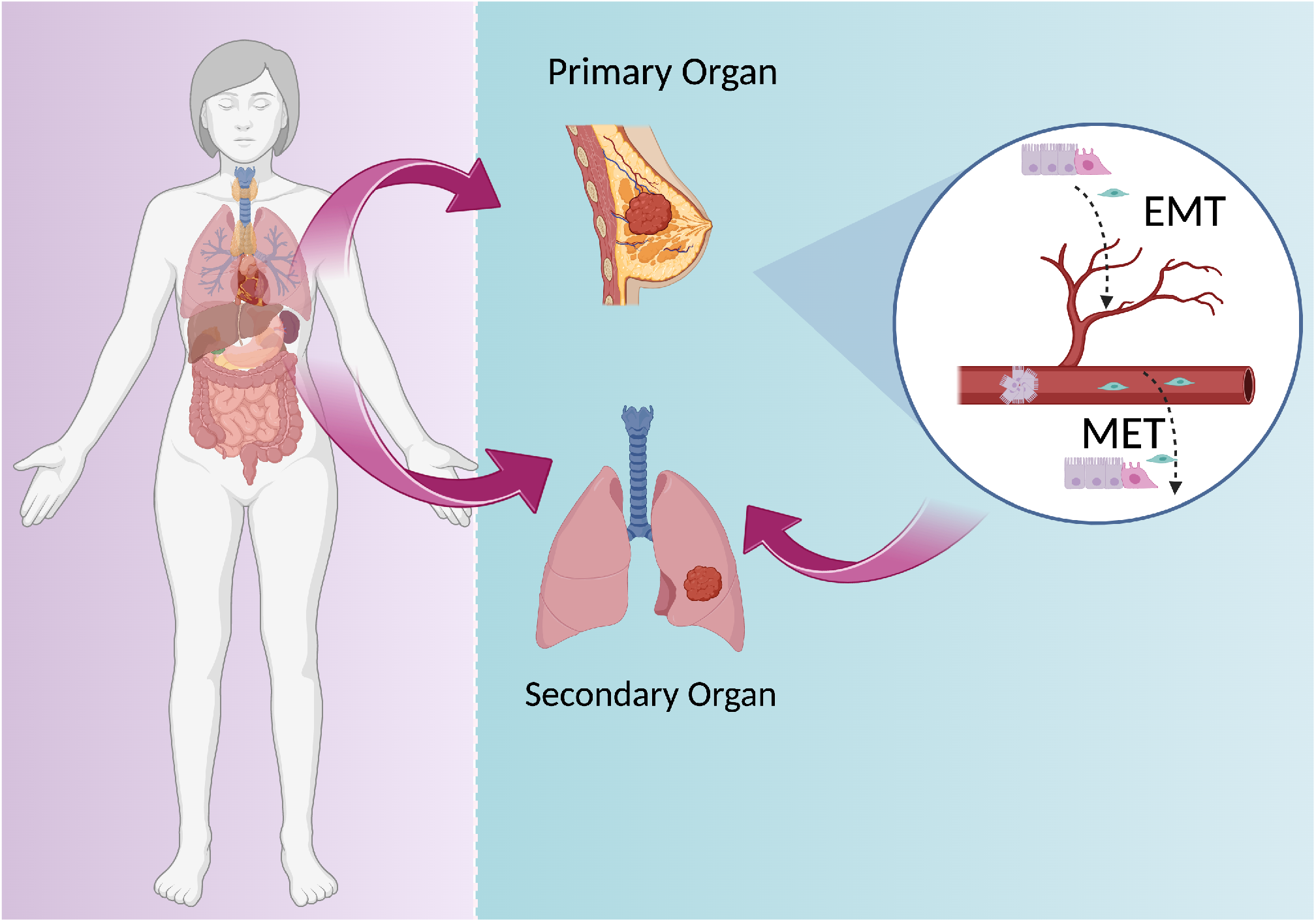

**Highlights:**

- A fully stochastic model elucidates the population dynamics of EMT
- A data-driven pipeline is introduced to track EMT trajectories from single-cell RNA sequencing
- Cell cycle scoring reveals cell line-dependent patterns of EMT Induction

## 1. Introduction

The Epithelial-Mesenchymal Transition (EMT) is a reversible dynamic cellular process involving the transformation of membrane-bound epithelial cells to motile mesenchymal cells. Although EMT was initially discovered in the setting of embryogenesis^57^, its importance in wound healing and the migration of cells to distant organs during metastasis is now appreciated and these topics have become active areas of research^19,37^.

EMT is driven by biomechanical and biochemical signals that lead to the loss of apical-basal polarity and cellcell junctions. These structural disturbances in organization arise due to the down-regulation of epithelial biomarkers and up-regulation of mesenchymal biomarkers^49,26^. Multiple EMT-associated transcription factors orchestrate EMT in an elaborate gene regulatory circuit^12,20,49^. As a result, EMT is not only affected by genetic alterations but is also driven by epigenetic and post-transcriptional modifications^53,38^. Thus, this process and its phenotypic implications are imperfectly understood by studying whole genome sequencing or allelotyping alone, particularly evident in studies attempting to identify mutant genes underlying metastasis ^64^

Studying EMT and its role in cancer progression is further complicated by the presence of non-cancer cells in the Tumor Microenvironment (TME), such as Cancer-Associated Fibroblasts^68,42^ and Tumor-Associated Macrophages which interact with and are in close proximity to tumor cells, leading to difficulties in distinguishing tumor cells en route to EMT from the TME^15^. Consequently, there is an ongoing debate on the exact contributions of EMT in cancer metastasis^14^: Mesenchymal cells are known to escape from the adaptive immune system, actively present in the tumor stroma^56,59^. On the other hand, epithelial cells can also break away from the tumor and travel collectively as circulating tumor cells (CTC) such that their epithelial marker expression patterns remain intact. While some studies report cells going through partial EMT, others report that the leaders at the edge of the clusters display promiscuous gene expression patterns while the inner layers remain more uniform^67,7^.

EMT conceptualization has benefited significantly from the theoretical prediction of a hybrid intermediate state whose properties fall on a spectrum between epithelial and mesenchymal phenotypes^32^. This phenotypic spectrum permits fine-tuned adaptations to environmental cues through cellular plasticity that can manifest as enhanced tumor aggressiveness in the clinic. Consequently, recent investigations have focused on the partial EMT state and its subsequent transition as the main culprit behind metastasis ^23,2^. Moreover, it is now appreciated that this hybrid state is non-transient and reinforced by phenotypic stability factors (PSF), such as GRHL2 and NFATc, which enhance tumor-initiation ^40,50^.

TGF*β* is a critical EMT inducer, both in-vitro and in-vivo, that acts through transcriptional regulation resulting in the down-regulation of E-cadherin (E-cad) and further upregulation of TGF*β* in a positive feedback loop^20^. TGF*β* EMT induction in-vitro is known to be reversible whereby TGF*β* withdrawal results in a reverse MET. Notably, not all cells are responsive to the EMT-inducing signal resulting in considerable phenotypic intra-tumoral heterogeneity and coexistence amongst multiple states ^66^ (Figure 1B). This has been reinforced by experimental observations supporting the existence of phenotypic transitions amongst EMT states as well as model-driven predictions of phenotypic heterogeneity generated by noisy cell division^21,58^. Furthermore, the environmental cues that govern these cell state transitions are highly context-specific^43,62^. This in turn highlights the importance of computational frameworks that interrogate context-specific EMT in the absence of cell-division.

**Figure 1:**
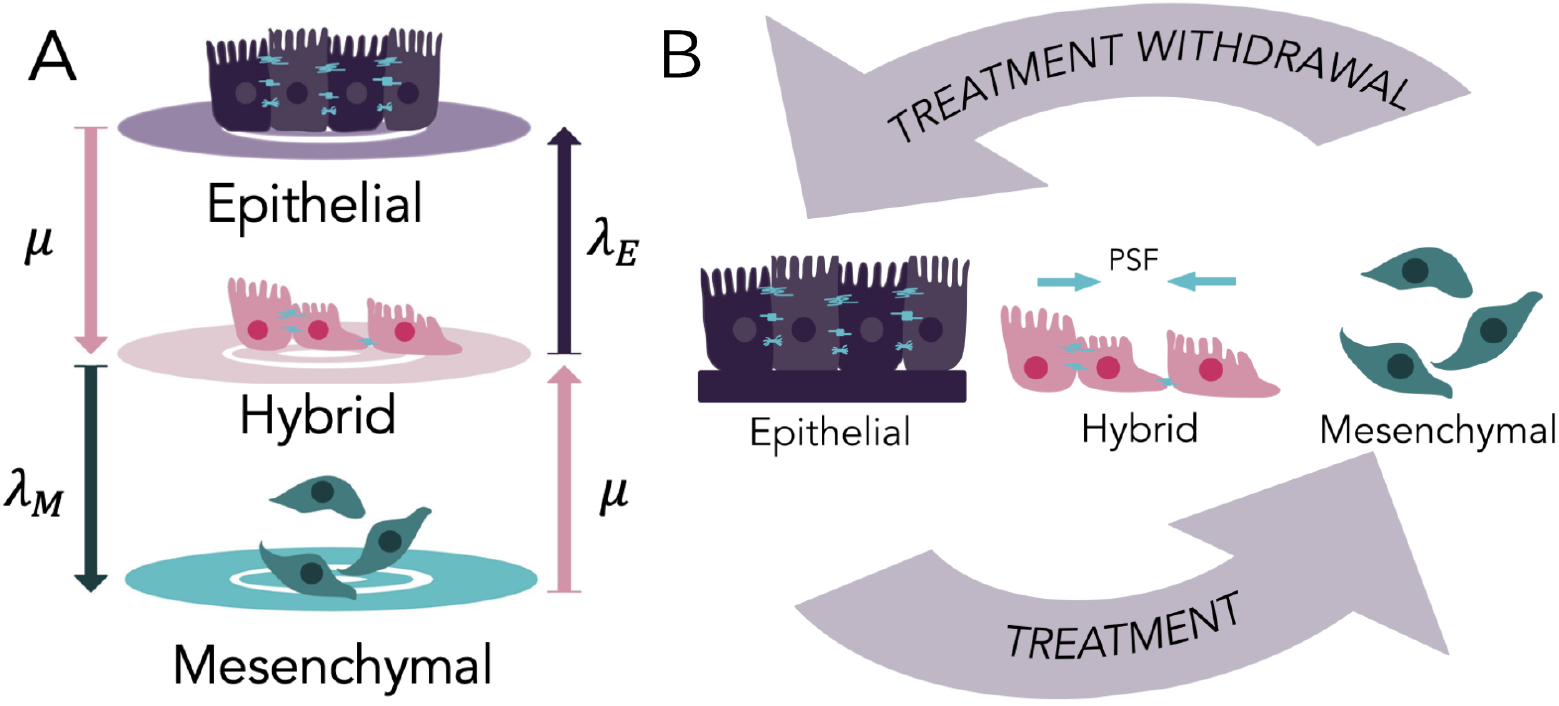
Stochastic Phenotypic Transition Model. (A) Illustration of the continuous-time three-state model and the four allowable transitions with their corresponding rates. Phenotypic stability factors enhance the stability of the intermediate state, and their corresponding transition rates (*μ*) are assumed to be symmetric.(B) Illustration of cell populations undergoing division-independent induced EMT, characterized by epithelial cells that lose their cell-cell junctions and apical-basal polarity. Upon treatment withdrawal cells regain their epithelial characteristics through MET. PSFs stabilize the hybrid state by making the transition rates into the hybrid state symmetric.

Prior computational frameworks have successfully inferred epithelial, hybrid, and mesenchymal states from transcriptomic data^16,54,3,4^. However, these methods were all optimized to explain microarray data, and are often ill-equipped to perform well on more modern approaches. Thus, there is a significant need to identify and predict context-specific EMT-related trajectories at transient and steady states from next-generation sequencing data.

Here, we provide a dual theoretical and data-driven framework, COMET (Cell line-specific Optimization Method of EMT Trajectories) for understanding the stochastic progression of EMT at stationary populations. We track the dynamics of these systems, which are treated with an EMT-induction factor, and show that COMET can successfully predict the timing and distribution of EMT states. Next, we accurately infer the three epithelial, hybrid, and mesenchymal trajectories from single-cell RNA sequencing data using COMET. We show that COMET explains early and late transition dynamics of context-specific EMT and relates the timing and equilibrium distribution to systemic noise through EMT induction exposure time. Additionally, we show that COMET reveals tumor subtype plays a more significant role than induction factors with respect to the dynamics of time-course EMT data which can give us insights into the patterns and extent of context-specific metastasis.

## 2. Model Development

### 2.1. EMT Population dynamics modeled with a three-state continuous-time Markov chain

We consider a non-dividing population of *N* cells, each of which belonging to one of three states; Epithelial (*E*), Hybrid (*H*), and Mesenchymal (*M*). We denote by *π_k_*(*t*) the relative abundance of population *k* (*k* ∈ {*E, H, M*}). For each time-point *t*, we have:

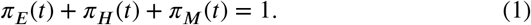

We model this process as a Continuous-Time Markov chain (CTMC)^27^. Letting *P*(*t*) denote the probability transition matrix of this Markov chain, this process evolves temporally via Equation 2:

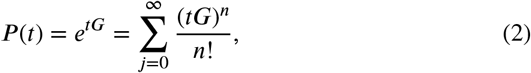

where *G* is the infinitesimal generator matrix representing the rates of transitions between the states of the system, given by Equation 3:

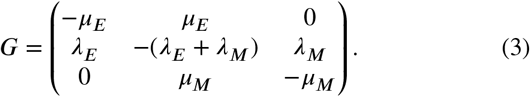

The rates *μ_k_* (resp. *λ_k_*) represent the transition rates out of (resp. into) state *k* (*k* ∈ {*E, M*}). It can be shown that this system permits a unique equilibrium distribution, which is represented by *π*_∞_ = [*π*_*E*,∞_, *π*_*H*, ∞_, *π*_*M*, ∞_] and satisfies

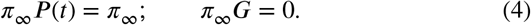

The stationary distribution in this case can be solved usings Eqs. 3–4 and is given by

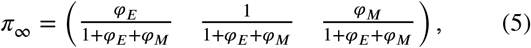

where *φ_i_* ≡ *λ_i_*/*μ_i_*.

### 2.2. Phenotypic stability factors augment transitions into the hybrid state

PSFs have been previously reported^50,40^ and act to stabilize the *H* phenotype. Here, we assume that their effect on EMT symmetrically enhances transitions into the *H* state (*μ_E_*=*μ_M_*=*μ* as shown in Figure 1A). In this case, we can explicitly derive the transition probability matrix, *P*(*t*), through eigenvalue decomposition of the generator matrix, *G* and Equation 2 (See Supplementary Information for detailed solution). *P*(*t*) can be represented a function of the stationary distribution and given by (Equation 6):

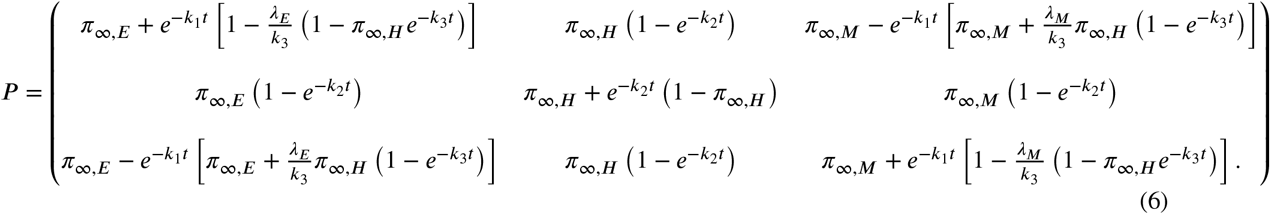

The temporal evolution described above can be extended by adding a spatial dimension. The dynamics of transitions in this case can be tracked by assuming a 1-dimensional signaling factor (such as TGF*β*) inversely influences the rates of transitions into the epithelial, and mesenchymal states (*λ_E_*, and *λ_M_*) through a sigmoidal function. We show in the Supplementary Information that in this case the hybrid state is constant in space (See Figure SI.1).

Additionally, the environment may be subject to discrete temporal changes. We let the times associated with each transition consist of an ordered set 0 = *T*_0_ < *T*_1_ < ··· < *T_i_* < … *T_M_*(*T_M_* denotes the terminal time), and denote by Δ*T_j_* ≡ *T_j_* – *T*_*j*–1_. We also denote by *λ_E,i_, λ_M,j_*, and *μ_i_* the rates associated on the interval *T_i_* ≤ *t_i_* ≤ *T*_*i*+1_. Then, for the stochastic matrix *P_i_* corresponding to *t_i_* we have via the Chapman-Kolmogorov Equation that,

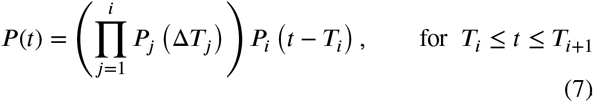

Our approach therefore can capably handle any description of temporal dynamics that may be divided into finitely many time-homogeneous regimes.

### 2.3. Symmetric noise tunes the temporal evolution of EMT trajectories

Empirical evidence supports a relationship between longer exposure time to the EMT stimulating signal and noisier gene expression patterns resulting from the stochasticity of the underlying biological mechanisms (intrinsic noise)^70^. Additionally, cells are subject to stochasticity in the stimulating signal resulting from longer exposure to the inducing factor (extrinsic noise)^30^. We account for both sources of noise using a *noise parameter α* that symmetrically scales the three transition rates such that 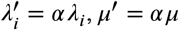. The *α* parameter influences the temporal evolution of the process without affecting the stationary distribution given in Equation 5.

## 3. Results

### 3.1. TGF*β* time-course flow cytometry data reveals early and late EMT dynamics

Our model was trained on multi-replicate flow cytometry data from Jia et al^22^. Cells were treated with 5 ng/mL of TGF*β* for a variable amount of time, after which treatment was withdrawn. This was repeated where withdrawal occurred for 3 biological replicates starting from day 3 to day 27 in increments of 3 days. The data was classified into four cases of RFP+GFP-(*E*), RFP+GFP+ (*H*), RFP-GFP+ (*M*), and RFP-GFP-. To isolate the three EMT-related trajectories, we normalized the three EMT-related cell fractions by the sum of *E*, *H*, and *M* cells.

In reconstructing the temporal evolution of the three replicates via the above processed signals, we noted a pattern of short-term phenotypic stability followed by a long-term transition to a steady state under continued EMT induction (Figure 2). More specifically, the temporal EMT distribution was characterized by an initial predominance of the *E* fraction that declined and transiently transitioned into a *H* phenotype as shown in Figure 2A and B. This *H* phenotype then slowly transitioned into a *M* phenotype resulting in the stable co-existence of *H* and *M* cells (Figure 2B and C). Surprisingly, the *H* and *M* trajectories of the data switched at day 18. However, in absence of any specific underlying biological reason, we considered this switch as an empirical artifact and discarded it for any training purposes. Motivated by these findings, we proceeded to fit a dual-regime CTMC to the time-course data.

**Figure 2:**
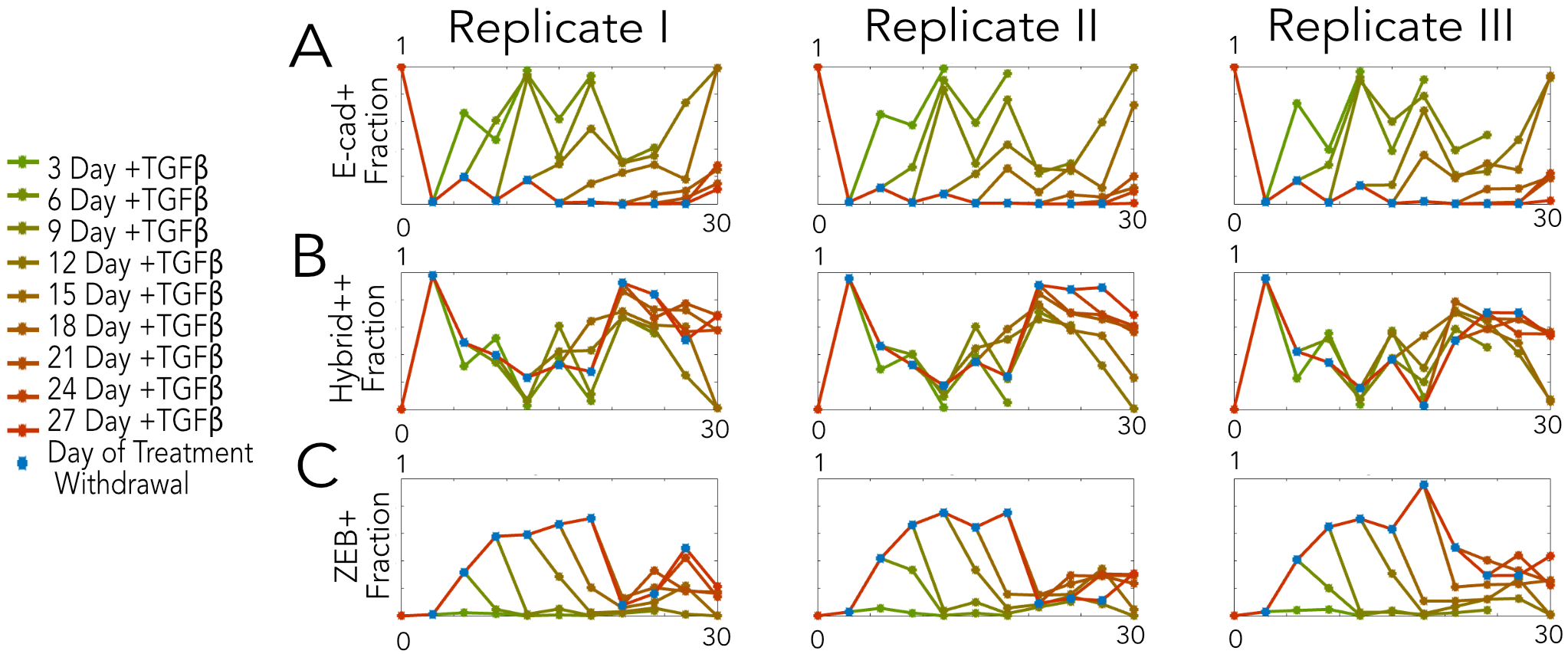
Timecourse Flow Cytometry Data. The experimental fraction of cells in (A) *E*, (B) *H*, and (C) *M* phenotypic states in time. The original flow cytometry data was normalized by omitting RFP-GFP-cells for each replicate^11^. Data shows a transient increase in the fraction of cells in the *H* state in the short-term that is followed by a transition into the *M* state long-term.

After calculating the mean of the trajectories over the three replicates, the first Markov chain was fitted from day 0 to the day where the *H* trajectory peaked (day 3). The system evolved via a probability transition matrix with parameters determined from an assumed hidden stationary distribution at day 3. We hypothesized that upon reaching that threshold the system switches to a different regime until relaxation to a steady state. The second regime then began at day 3 and relaxed to steady state with the distribution at day 18 assumed to be the stationary distribution. This regime was normalized in time relative to the first regime, which was selected arbitrarily (we further discuss the approximation steady-state using day 18 values in Section 3.3).

Increases in the relaxation time to equilibrium distributions observed during the second regime were accounted for by applying the noise parameter discussed above to optimally fit the theoretic trajectories to empirical data in this regime (Figure 3A). Without loss of generality, we first set one of the transition rates (*λ_E_*) to *α* and solved for the others by equating Equation 5 to the empirical stationary distribution. Optimization was performed using the *fminsearch* function in Matlab to find the optimal *α* minimizing the total squared error between the three theoretic and corresponding empirical trajectories^29^ (Figure 3B). For further validation, we also applied a non-linear unconstrained optimization method, *fminunc*, that utilizes gradient descent and confirmed agreement in optimal parameters (See Figure SI.2). Collectively, we find that our two-regime approach performs well at characterizing empirical phenotypic fractions.

**Figure 3:**
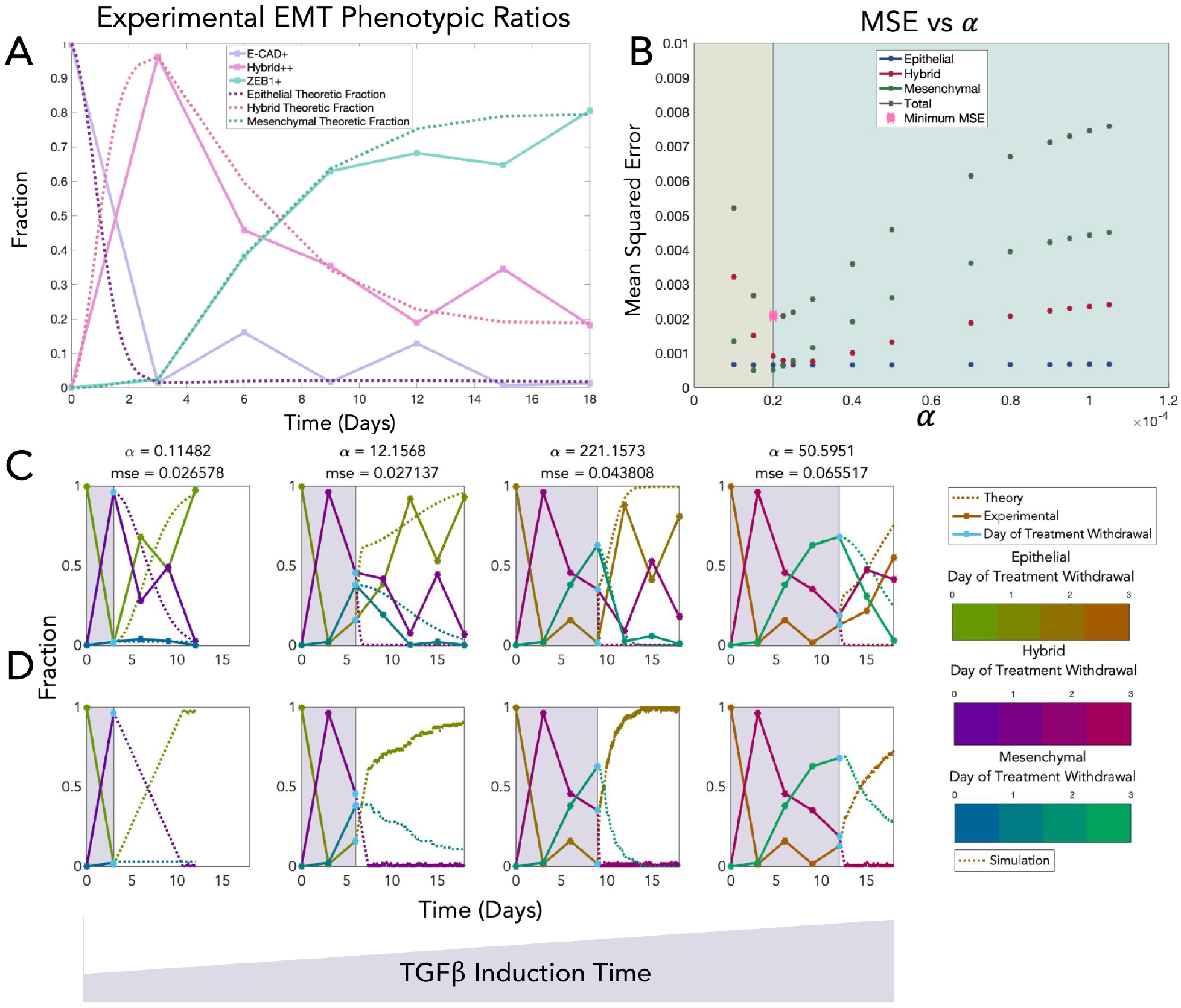
Stochastic Modeling Framework Applied to Time-course EMT Data. (A) The two-regime Markov chain (dotted) fits the non-monotonic phenotypic composition of EMT induction observed empirically (solid, averaged over 3 biological replicates). (B) Numerical optimization is performed to fit the time-course data in part (A) by identifying the best-fit *α*. (C) Experimental and theoretical phenotypic trajectories for EMT through time as a function of TGF*β* treatment induction explained by increasing values of fitted *α*(D) This panel shows how stochastic Gillespie simulations generate distributions that are in large agreement with the experimental results. Gillespie simulations were performed over sufficiently large number of iterations to ensure steady state was reached (10000). The resulting values were then trimmed and normalized based on their inter-arrival times. The smallest inter-arrival times belonged to the third case (treatment withdrawal at day 9). This panel shows only one run of the simulations.

### 3.2. COMET’s CTMC framework predicts enhanced transition rates en route to MET as a result of longer TGF*β* exposure

The same procedure was repeated to minimize the Mean Squared Error (MSE) between the three theoretic and corresponding empirical trajectories following TGF*β* treatment withdrawal. Here, we assumed that phenotypic reversion would ultimately recapitulate the initial, pre-TGF*β* distributions. Using this approach, we found that the *α* values correlate directly with TGF*β* treatment exposure (Figure 3C). In addition, we simulated the random arrival of the four possible transitions between the three states as shown in Figure 3D, using Gillespie algorithm^17^ and found great consistency between simulations and theoretic fractions.

### 3.3. COMET infers the three EMT-related trajectories from single-cell RNA sequencing data

To evaluate our theoretical CTMC model on additional empirical data and to extend our framework into an analytical tool, we developed a data-driven pipeline that enables inference of EMT-related trajectories from single-cell RNA sequencing data. Time-dependent data of four cell lines (A549, DU145, MCF7, and OVCA420) treated with three EMT induction factors (10 ng/ml of TGF*β*, 10 ng/ml of TNF, and 30 ng/ml EGF)^9^, and dose-dependent steady state data of MCF10A cell line treated with various doses of TGF*β* ^41^ were used to infer the three EMT-related trajectories.

We applied initial quality control (which included filtering based on the number of expressed housekeeping genes, total expressed genes, and mitochondrial percentage), followed by library size normalization and data filtration using previously reported EMT-related genes^16^. Next, we extracted the top 100 most variable genes using Seurat^46^ in R across each available combination of cell line and induction factor^9^. We found that the top variable genes were largely shared among cell lines rather than EMT induction factors (Figure 4A).

**Figure 4:**
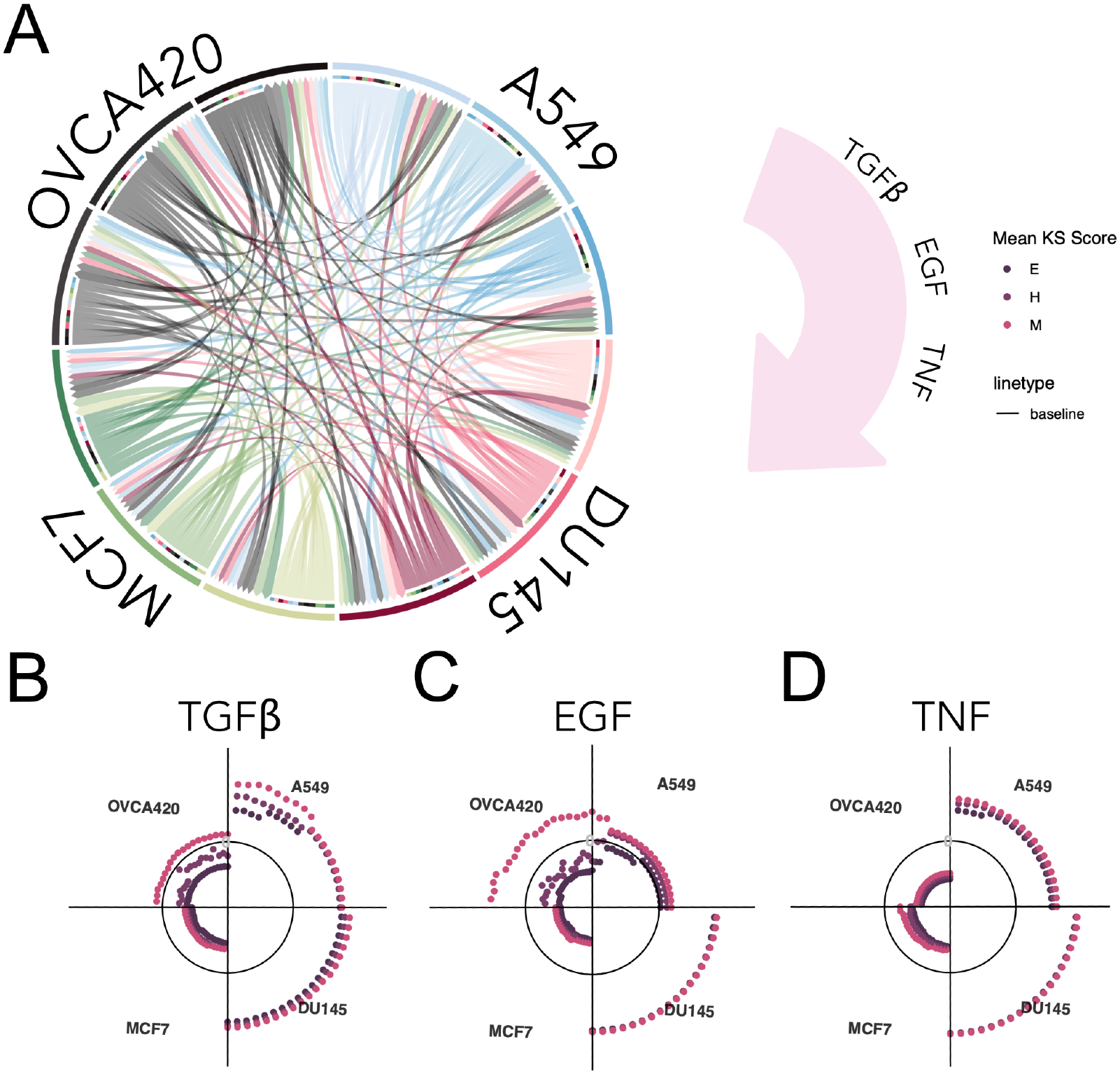
Data-driven Pipeline Scoring of Time-dependent Data. (A) Shows the Circos plot for the top 100 highly variable genes. The plot demonstrates that highly variable EMT-associated genes are shared among cell lines rather than across treatments. The length of the incoming arrows to every subsection of the plot indicates the number of genes that are common between two cases. Each colored slice is related to a specific induction factor (clockwise lighter to darker: TGF*β*, EGF, and TNF) (B-D) Each panel slice represents the mean KS score over 10 runs of the algorithm when using 10 to 100 highly variable genes (increasing clockwise in increments of 5). As shown in the plots, the mean KS scores appear to follow the same pattern across cell lines and do not depend on the EMT induction factors.

We then filtered the data further by identifying the 10 most highly variable genes, followed by imputation via MAGIC^61^. This was followed by dimensionality reduction via UMAP ^34^ and K-means clustering (The UMAP plot for one run of the algorithm - with data filtered for the optimal number of EMT genes - for the four cell lines is shown in Figure 5A). We then labeled the three clusters using a previously developed EMT metric, the Kolmogorov-Smirnov (KS) based method^54^. KS scoring was performed on every cell based on their MAGIC imputed genetic profile with respect to the top 100 variable genes included. This decision was made to ensure enough genes exist for KS scoring. Lastly, in order to associate the three clusters with *E*, *H*, and *M* phenotypes, we took the average of the KS scores for every cluster and sorted the clusters such that low, intermediate, and high KS scores corresponded with *E*, *H* and *M* states, respectively.

**Figure 5:**
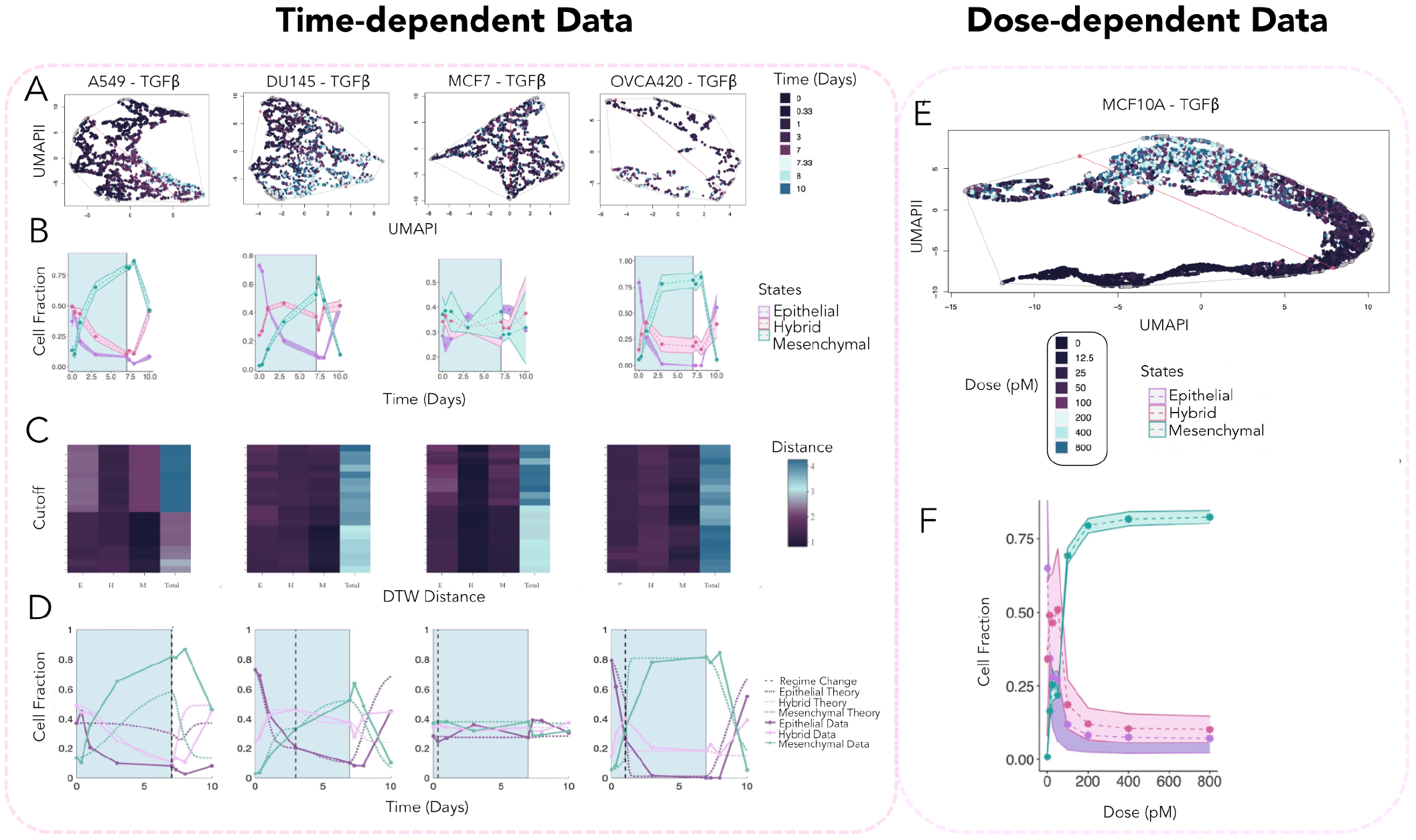
COMET Predicts the Timing and Distribution of Context-specific EMT. Figures (A), (B), (C), and (D) are timedependent data of four cell lines treated with 10 ng/ml of TGF*β* for seven days which underwent MET following treatment withdrawal for three days. Figures (E) and (F) represent the steady state information of the MCF10A cell line treated with various doses of TGF*β*. (A) Shows the UMAP plots for the four cell lines treated with TGF*β*. Archetypal analysis with two archetypes^13^ was performed on data and is shown in the figure. The figures demonstrate a clear transition from the *E* archetype to the *M* archetype for the three cases of A549, DU145, and OVCA420. (B) This panel shows the inferred time-course trajectories with confidence intervals for 10 runs of the algorithm. The blue region depicts the duration of treatment. (C) Heatmaps show the DTW distances resulting from the DTW alignment of the three EMT trajectories inferred using the pipeline and flow cytometry data. The lowest total DTW distance was used to choose the cutoff of highly variable genes which resulted in the time-course trajectories in the panel above. (D) The stochastic model was fitted to the mean trajectories of the 10 runs for the optimal cutoff. (E) The UMAP plot shows a transition from the *E* to *M* phenotypes as a function of treatment dose (gene cutoff 45). (F) Plot illustrates the inferred trajectories from the data-driven pipeline for the dose-dependent data. The specific transition rates for each case are reported in Figure SI.25.

From the fraction of cells in every cluster we inferred the three EMT-related trajectories and repeated the same procedure as before, this time varying the number of included variable genes starting from the top 15 to the top 100 in increments of 5. Due to the stochasticity of our algorithm, we repeated this procedure 10 times for each cutoff and found the mean KS scores (Figure 4B-D.).

Consistent with the original study and other investigations of this data^44,9^, the pattern of the mean KS scores was found to be highly dependent on the cell line than EMT induction factor (See Supplementary Information for reported KS scores). This finding, in light of the conservation of variable genes within cell lines experiencing distinct induction treatment, further substantiates our prior finding that phenotypic transitions are dominated largely by the particular cell line, and not the particular induction factor, involved in undergoing an EMT. Our approach predicts that A549 and DU145 cell lines exhibit higher mean KS scores (A549-TGF*β*: 0.501, DU145-TGF*β*: 0.559, A549-EGF: 0.059, DU145-EGF: 0.596, A549-TNF: 0.405, and DU145-TNF: 0.648) compared to OVCA420 and MCF7 (OVCA420-TGF*β*: −0.110, MCF7-TGF*β*: −0.276, OVCA420-EGF: −0.074, MCF7-EGF: −0.340, OVCA420-TNF: –0.374, and MCF7-TNF: −0.259) over a majority of cutoffs (Figure 4B-D), which is consistent with previously reported findings of MCF7^16^ and OVCA420^51^ exhibiting *E* characteristics. Moreover, prior work identified a higher percentage of hybrid cells for DU145 and coexistence *E*, *H*, and *M* fractions for A549^16^, which can also be appreciated by constructing our inferred trajectories for day 0 (Figure 5B). Intriguingly, this suggests that our approach reliably quantifies EMT phenotypic composition at the single-cell resolution from next generation sequencing data.

Next, to account for the variability in different runs the algorithm, we calculated confidence intervals for every trajectory (See Figures SI.8-19 for inferred trajectories with confidence intervals. Figure 5A, and B show the UMAP and inferred trajectories with confidence intervals for the optimal cutoffs of highly variable EMT genes respectively) Occasionally, we observed that the inclusion of 5 additional genes drastically changes the EMT trajectories (Figures SI.8-19). This is likely due to an abrupt change in the number of resolvable *H* states based on the genes included which could not be detected by our three-state model. Our pipeline inferred ambiguous and highly variable trajectories for the majority of the EGF and TNF cohort (all inferred trajectories for these cases are reported in Figures SI.12-19). We note that from the phase contrast images reported by Cook and Vanderhyden^9^, it is indeed unclear whether cells went through EMT in these cases. As a result, we proceeded with our subsequent analysis performed on TGF*β*-treatec cell lines only.

We hypothesized that EMT in this additional dataset would temporally evolve in a similar fashion as the flow cytometry data of Jia et al. ^22^ and could thus be explained by our framework. Toward this end, we optimized the number of EMT-related genes that result in mean trajectories most similar to those of the originally considered flow cytometry data by utilizing Dynamic Time Warping (DTW) alignment DTW alignment optimizes the alignment of time-course trajectories by tweaking the time axis recursively^47^. As a result, the DTW distance can be used to measure similarities in dynamics of transitions by ignoring the context-specific timing of events.

We performed DTW alignment of the single-cell RNA sequencing trajectories with the corresponding trajectories of the mean signal of the flow cytometry data. The resulting heatmap which illustrates the DTW distances for every cutoff of highly variable genes is shown in Figure 5C. The cutoff with the lowest sum of DTW distances of the three trajectories from the flow cytometry data was considered for fitting the stochastic model to time-dependent data. While for the optimal cases we observed cells to be on a spectrum from epithelial to mesenchymal phenotypes on the UMAP plot (Figures SI.3-6), we noticed that the measured EMT spectrum visualized by UMAP plots became less discernible as we increased or decreased the number of included highly variable genes (Figures SI.3-6). To quantify this phenomenon, we fit a minimum spanning tree (MST) to the UMAP plot to measure the pairwise distances between cells in Figure SI.7 and showed that the maximum edge of the tree is minimized in the neighborhood of the optimal cutoff of variable gene (the minima fall between a cutoff of 40-55 for all TGF*β* cases; Figure SI.7).

We next proceeded to test our results on an independent dataset featuring dose-dependent EMT induction^41^. Since the optimal cutoff of highly variable genes fell uniformly between 40-55, we considered a cutoff of the top 45 highly variable genes for this analysis (Figure 5E depicts the dimension reduction plot for the dose-dependent data where cells treated with low doses of TGF*β* at steady state move from one cluster in the neighborhood of an *E* archetype to an *M* phenotype as the dose of treatment increases). The resulting optimal trajectories for the time-dependent and dose-dependent data with confidence intervals are shown in Figure 5B and 5F respectively. Although our 45-gene approach exhibited some differences between the predicted and simulated phenotypic fractions as a function of dose reported by previous studies, our overall trends were in general agreement^35^ (Figure 5F). From the steady state fractions obtained from the dose-dependent data, it is evident that the distribution of fractions for the flow cytometry data at day 18 oesemoles the equilibrium state fractions closely (roughly 113.6 pM, see Supplementary Information for dimensionality analysis).

### 3.4. COMET CTMC predicts time dynamics of context-specific EMT

To further evaluate the dynamical effects of various induction factors on a cell-line-specific basis, we next fitted COMET’s CTMC model to time-dependent data of each of the four cell lines treated with TGF*β*. We then fitted Markov chains from day 0 to the day where the *H* state peaked with the timescale normalized based on the flow cytometry data. Given our prior analysis on empirical time-course proportions, we performed this normalization assuming that TGF*β* induction, with a different dose on a distinct cell line, would follow similar dynamics en route to EMT. However, the stationary distribution of both regimes was determined based on the time-dependent data due to the discrepancies in cell line type and dose of TGF*β* as well as lack of available steady state information. For one of the cell lines, A549, the *H* state was maximized at day 0 thus we only fitted a single Markov chain to the data, assuming this case starts within the second regime. This assumption is supported by the fact that CEACAM6, an inducer of EMT^6^, is the most variably expressed gene at day 0 for A549 across all treatment cases (Figures SI.20-22). For all cases, the second regime of the Markov chain was fitted from the day when the hybrid state peaked to the day of treatment withdrawal (day 7).

Next, we proceeded to apply the noise parameter, *α*, to explain changes in the temporal evolution of the Markov chain. We then fitted a separate Markov chain following treatment withdrawal from day 7 to the end of the experiment (day 10) assuming the trajectories revert back to the initial distribution at day 0. Similarly the optimal fit was obtained using the *α* parameter. The final results are shown in Figure 5D. Since our optimization depends on the MSE of three trajectories, occasionally an optimized theoretic curve only fits one trajectory well while poorly fitting the other two (A549 of Figure 5D). In these cases a similar MSE can be obtained through a larger or smaller *α* parameter that optimally fits the other two trajectories. Overall, we observed reasonable consistency in our inferred trajectories and the CTMC theoretic trajectories (See Supplementary Information and Figure SI.25 for the exact inter-state transition rates).

### 3.5. Cell cycle scoring of the time-dependent data reveals cell line-dependent cell cycle fractions

To further evaluate the utility of COMET, we next interrogated the growth rates of cells by extracting cell cycle fractions through Seurat normalization and performing cell cycle scoring^46^. Consistent with previous reports^65^, we found cell cycle stages to be similar across different cell lines for the time-dependent data as shown in Figures 6B-D.

**Figure 6:**
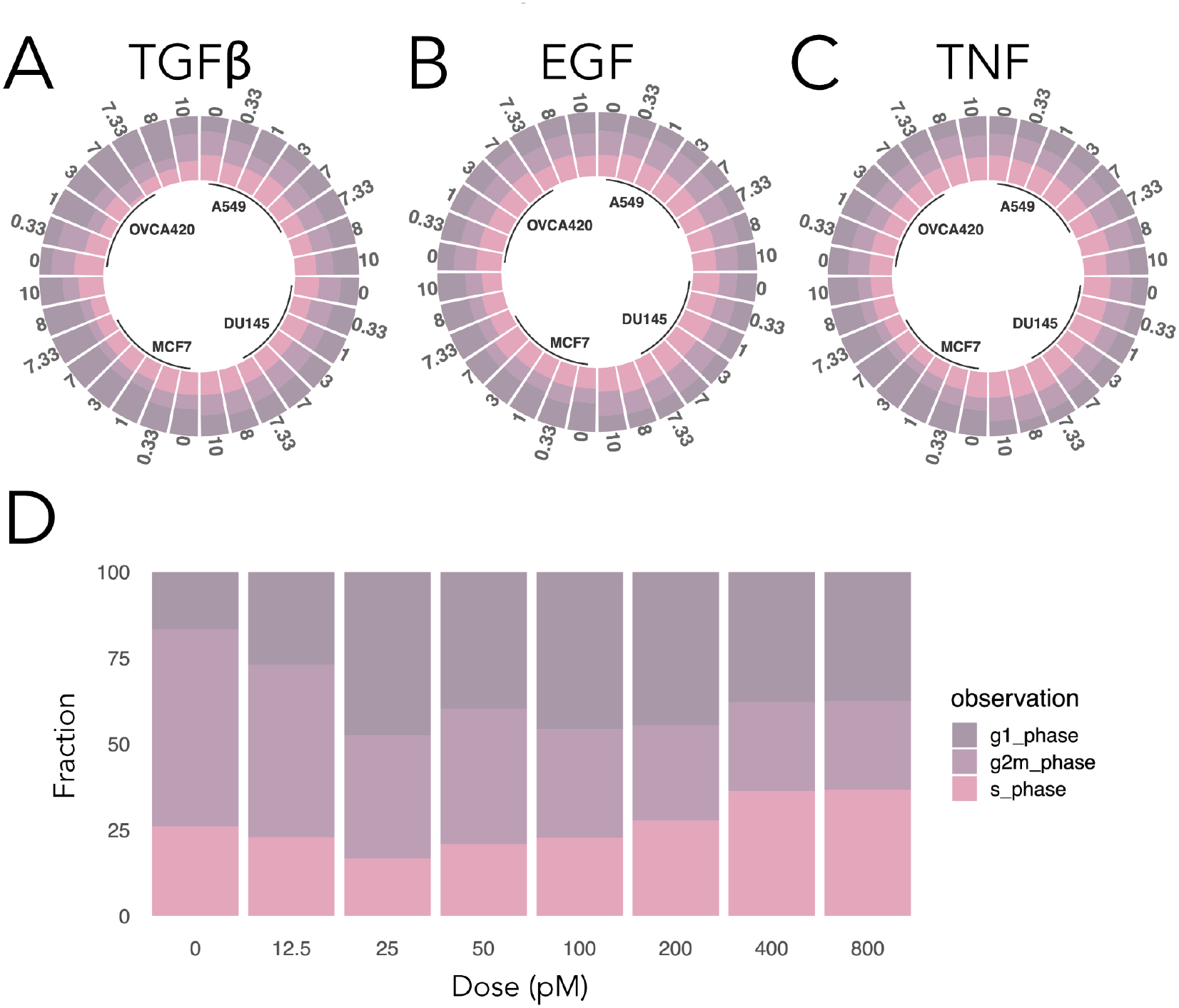
Time-dependent and Dose-dependent Data Cell Cycle Scoring. (A-C) Illustrate the cell cycle scores for every cell line of the time-dependent data. Similar to the KS scores, cell cycle scores appear to be more cell line dependent than induction factor specific. (D) Shows the cell cycle fractions for the dose-dependent data. Cell cycle fractions at steady state are reported as a function of TGF*β* dose. Cell cycle informati n starts with a high percentage of cells in the G2M phase which declines, followed by a transient increase in the G1 phase and an increase in S phase long-term.

We assessed the trends in G1 phase of cell cycle stages of time-dependent data during TGF*β* treatment using Kendall Tau statistic^28^ and found that in general the G1 stage positively trends for TGF*β* treated cells during treatment (day 0 to 7) as shown in Figure 6A, consistent with previous reports ^52^. However, this trend was stronger for the two cell lines with lower mean KS scores (MCF7: 0.6, and OVCA420: 0.8) than DU145 and A549 (DU145: −0.2, and A549: 0.4) (Figure 6A). This is consistent with previous reports of normal epithelial cells arresting at G1 phase following TGF*β* EMT induction while those with high cell proliferation fail at cell cycle arrest resulting from genomic instabilities and mitotic defects^8^.

We also discovered a higher proportion of cells in S and G2M phases that happened to coincide with the two cell lines having higher KS scores (DU145, and A549) as they progressed through EMT. Furthermore, we found the fraction of cells in G1 phase to be relatively high for the MCF7 cell line compared to the other cell lines at baseline day. Surprisingly, in cell cycle scoring the dose-dependent data we observed the fraction of cells in the S phase increasing as a function of TGF*β* dose (Figure 6D). Consistent with studies reporting an increase in the fraction of cells in S phase as a result of a decrease in p21 expression while cells go through EMT^63^, COMET predicted an increase in the fraction of *M* cells as a function of dose for the optimal cutoff (Figure 5F). However, as reported in Figure SI.24, for a cutoff of 100 highly variable genes COMET predicted a higher percentage of cells in the hybrid state for the dose-dependent data.

### 3.6. GSEA resolves enrichment in cancer-specific hallmarks for each of the EMT-related phenotypes across distinct cell lines

Lastly, to further test the ability of COMET to accurately infer the three EMT states from time-dependent single-cell RNA sequencing data, we performed Gene Set Enrichment Analysis (GSEA) on every run of the pipeline for the optimal number of highly variable genes. We start our analysis with the TGF*β* signaling pathway since we assume cells would be up-regulated for the TGF*β* signaling pathway when undergoing EMT and downregulated following treatment withdrawal with the intensity of these patterns dependent on the rate and fraction of cells undergoing each process.

For instance, as shown in Figure 5D, a large fraction of *E* cells in the A549 sample appear to have undergone EMT prior to TGF*β* treatment, appearing stationary posttreatment. This would suggest that the *E* cluster may be significantly enriched for TGF*β* signaling, which we in fact confirm (Figure 7C). Similarly, the dynamics of the *M* proportion appear to indicate a rapid decline following treatment withdrawal and indeed we find that the TGF*β* signaling pathway is depleted in this case. Lastly, the *H* trajectory of the A549 cell line appears to have a mixed response where TGF*β* withdrawal results in more noticeable effects than induction.

**Figure 7:**
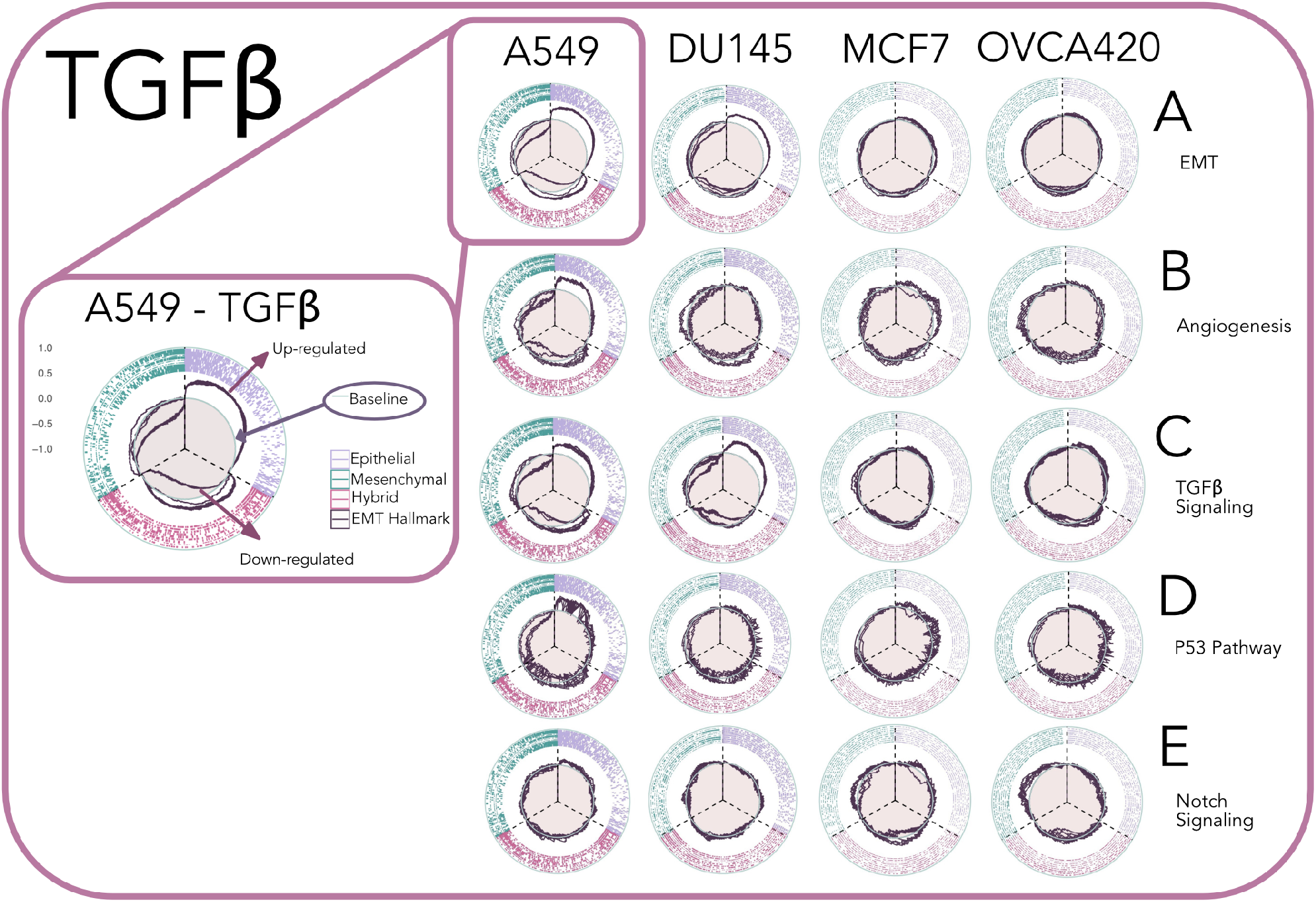
GSEA Plots of Relevant Hallmarks for Each Phenotypic State. GSEA plots are overlapped for 10 runs of the algorithm. The baseline separates the up-regulated and down-regulated genes. Every one-third of the circular plot bound by dashed lines represents one phenotypic state (Lines falling inside the baseline circle demonstrate an abundance of down-regulated hallmark genes and lines falling outside the circle show an abundance of up-regulated hallmark genes). (A) Shows cluster-specific GSEA plot for the EMT hallmarks, it is evident from the plot that the EMT hallmark gene set is more variable across states for DU145 and A549. An overall up-regulation trend for the EMT gene set transitions to a down-regulation trend for the mesenchymal state. GSEA plots for the three phenotypic states are given for (B) angiogenesis, (C) TGF*β* signaling, (D) P53 pathway, and (E) notch signaling.

A similar pattern is observed For DU145. In this case, the majority of the *E* cells reside in the pre-treatment regime thus significant up-regulation of the TGF*β* signaling pathway occurs for the *E* cluster. While the *M* fraction almost reverts back to its initial fraction at a shorter period. Thus, TGF*β* signaling pathway is down-regulated for the *M* trajectory. On the other hand, the *H* fraction is almost stabilized across time and since the other cells are transitioning into the *H* state, a larger fraction of hybrid cells are up-regulated for TGF*β* pre-treatment.

In contrast, for the MCF7 case, the three trajectories appear to stably coexist and thus we do not observe either up-regulation or down-regulation for any of the clusters. As mentioned previously, we are unsure whether this cell line underwent EMT in the original data^9^. Lastly, for the OVCA420 cell line almost all of the three trajectories revert back to their initial fraction and the pattern of EMT progression and MET is fairly symmetric. The GSEA analysis faithfully captures this observation, with enrichment scores approaching zero with small deviance for the *E* and *H* trajectories (as they do not fully revert back to the initial day fractions).

We then analyzed the EMT hallmark cases and noted their similarity to the TGF*β* signaling pathway enrichment scores. However, issues arise during GSEA analysis of EMT-related clusters due to the concomitant existence of both *E* and *M* biomarkers in the gene set. As a result, net zero enrichment may be reported when, for example, the up-regulation of *E* markers happens concurrently with the down-regulation of *M* markers (Figure 7A).

We observed a transition from up-regulation to down-regulation from *E* to *M* clusters for the P53 pathway gene set across all cell lines (Figure 7D). This collective pattern is consistent with previous reports that highlight the role of the P53 pathway in regulating EMT by binding to the miR-200c promoter^5^. Surprisingly, this contrasted with our observation of different patterns of up-regulation or downregulation of the angiogenesis hallmarks across different cell lines. While angiogenesis followed the same pattern of the EMT gene set in the case of A549 (Figure 7A and B), it followed the exact opposite pattern for DU145 (Figure 7B). Although the tumor stroma is known to influence patterns of vascularization for different tumor types, our results suggest tumor sub-type-specific patterns of angiogenesis in the absence of tumor stroma and under the influence of the same signaling factor^33^.

From the gene set enrichment analysis of cell lines for the notch signaling pathway, we observed a pattern of up-regulation to down-regulation only for the OVCA420 case. For DU145 and MCF7, we did not observe significant up-regulation or down-regulation of the notch signaling pathway. However, for A549 we observed a slight up-regulation of the notch signaling pathway. This result is inconsistent with previous reports of TGF*β* directly resulting in the up-regulation of notch ligand after EMT induction. However, the cell lines used in this study are different and from Figure 7E, our results suggest tumor type dependent pattern for notch signaling activation^69^.

As shown in Figure 7A-D, the patterns of GSEA up-regulation or down-regulation are more pronounced across the four gene sets of EMT, angiogenesis, TGF*β* signa?.ig, and P53 pathway in the cell lines with higher KS scores (A549, and DU145) versus those with lower KS scores (OVCA420, and MCF7).

Furthermore, we note a similar pattern of GSEA enrichment for clusters across several different hallmarks for the same cell line. We evaluated the similarities between gene sets via pairwise Jaccard distance and the number of shared genes across gene sets fell below 1%. This result suggests extensive cross talks between these processes and the existence of gene regulatory networks controlling the concurrent up-regulation or down-regulation of gene sets across cell lines. Collectively, enrichment analysis further validates COMET’s characterization of single-cell RNASeq signatures of each of the three relevant phenotypes, and can be more generally used to investigate pathways that are actively involved in maintaining each state.

## 4. Discussion

Despite the vast body of EMT-related studies, our understanding of EMT and its specific role in metastasis is far from complete. Due to the multifaceted role of transcriptional regulation in driving EMT^55^, interrogating metastasis at the genetic level has often yielded in inaccurate findings^64^, and there is a larger need for tools to infer the dynamics of EMT at the phenotypic level.

Here, we presented an analytic tool, COMET, for reliably inferring and predicting EMT trajectories from single-cell RNA sequencing data. We applied COMET to time-course data of cell lines treated with TGF*β*, and showed that our method was able to recapitulate the findings of previous reports at the single-cell level^16^. Reliable identification of EMT-related phenotypic fractions from increasingly available single-cell RNA sequencing data will enable us to better understand EMT and its pattern of progression in cancer.

Using COMET, we showed that our inferred trajectories collectively followed a pattern of a short-term monotonic increase in the *H* state followed by a transition to the *M* phenotype leading to the stable coexistence of the two phenotypes. This pattern was exceptionally not observed for the MCF7 data among the TGF*β*-treated cohort. We note that based on the phase contrast images of the MCF7 cell line reported by Cook and Vanderhyden^9^, cells do not display fully epithelial characteristics at day 0 and their progression through EMT is subsequently unclear. As a result, this heterogeneity may confound the ability of our pipeline to resolve the EMT trajectories where the gene expression signature does not undergo phenotypic transitions^39^. However, this observation was constricted to the time length of the study. We note that for the single-cell RNA sequencing data, cell lines were only treated with the EMT induction factor for 7 days and as the original paper states it is unclear whether cells reached steady state distribution at day 7 ^9^.

Adding to this complexity is the fact that we observed a switch between the *H* and *M* trajectories at day 18 from the ‘ow cytometry data of Jia et al^22^ which was acquired over a longer period of 30 days. Although we discarded data from days 18 - 30, assuming the switch is an empirical artifact, this temporal inhomogeneity can be theoretically modeled via a separate CTMC regime. However, the lack of available gene expression data for this period limited our ability to investigate the underlying reasons behind this empirical switch through other concurrent processes such as cell cycle scoring or GSEA. Furthermore, this pattern of a switch between *H* and *M* phenotypes was not consistent with our dose-dependent steady-state distributions obtained from the data of Panchy et al^41^. Nonetheless, our analytic tool, COMET, provides researchers with the ability of reliably investigating this switch further on longer time-course single-cell RNA sequencing data in the future.

Our approach, while useful for robustly characterizing EMT at the single-cell level, is not without limitation. Phenotypically heterogeneous cells can exhibit gene expression profiles that cooperate or interfere with overall signal detection^1^, and this is further compounded by noisy celular division^21^. Although we have assumed in our analysis that populations under study were non-dividing, cell cycle scoring suggested the variable presence of division signatures across available experimental contexts as shown in Figure 6. However, despite this simplifying assumption, COMET performed exceptionally well in cases of even higher proliferating cell lines such as DU145 which has an abundance of cells in S phase (See Figure 6A). This result may suggest that the dominating contribution to EMT heterogeneity in the datasets considered here are driven by stochastic transitions rather than by cell-division.

Furthermore, there is a lack of consensus in the literature on the number of *H* phenotypes^24^ with their estimation possibly dependent on the number of biomarkers considered for EMT classification. Additional studies have also suggested that EMT is a non-Markovian process due to the existence of several microstates within macrostates^18^. In our model, we specifically optimized the number of EMT biomarkers based on their ability to resolve the data into three states depending on their sample-specific gene expression variation. Using a rigorous validation procedure, we showed that the number of highly variable genes, selected as representative of EMT and commonly chosen arbitrarily in computational studies, can drastically change the results of the analysis. Although our general findings on multiple independent time- and dose-dependent datasets^9,41^ were consistent with previous reports, due to the lack of predefined sample-specific *E*, *H*, and *M* fractions from single-cell RNA sequencing data, subsequent analysis on additional EMT-specific datasets will, further test COMET’s predictive accuracy. Our modeling approach, given sufficient availability of additional detailed data, could be used in an identical manner to account for and study multiple intermediate phenotypes in EMT.

Using COMET, our data-driven pipeline captured the phenotypic spectrum observed following dimension reduction of EMT-related genes and found through archetypal analysis with two archetypes (*E* and *M*) that the *H* state with an intermediate KS score is always spatially intermediary to *E* and *M*. This observation is in broad agreement with the recent specialist-generalist frameworks devised for the EMT process whereby the *E* and *M* populations were predicted to be optimized for one task at hand, while the *H* population confers fitness advantages owing to better collective performance at multiple tasks^10^.

In addition, previous models included the possibility of directly transitioning from an epithelial state into a mesenchymal state during EMT^25^, and many more report that cells transition into an epithelial state without ever visiting the intermediate state during MET^31,45^. In our approach, we decided to assume all transitions proceed through an intermediate state as cells are shown to retrace the EMT footsteps through continuum of positions in the space of EMT-related genes while going through MET. However, we note that this reverse transition happens much faster than EMT, thus it may be more difficult to observe this phenomenon empirically.

One of the main assumptions of our model was the symmetric effects of PSFs on the transition rates into the *H* state. This is supported to an extent by the symmetric trajecrories and consistency between predicted theoretic and inferred trajectories in Figure 5D. However, some evidence has suggested that the stronger presence of PSFs delays the transition into the *H* state in addition to the longer mean residence time within the *H* state^50^. We show through Gillespie simulation in Figure SI.23 that the short-term stability of the EMT-related trajectories can be explained by relaxing this assumption and the entire process can be modeled via a single CTMC (See Supplementary Information for full solution). Future work will further evaluate the performance of a single CTMC with asymmetric transition rates into the hybrid state.

Our analysis showed concurrent up-regulation and down-regulation of the angiogenesis, TGF*β* signaling, and P53 pathway hallmarks consistent with the EMT hallmark. The low number of common genes between these sets suggests the possible existence of a strong gene regulatory network controlling these processes. Future work inferring the gene regulatory networks governing context-specific EMT would provide additional insights into the underlying mechanisms that govern the phenotypic transition rates.

Lastly, to further extend our framework to in-vivo data at the tumor setting, we need to account for the difficulties of distinguishing stem-like cells from the elements of the TME and detecting them in circulation in-vivo^14^. Adding to this complexity is the fact that cells can also break away from the primary tumor and travel collectively as CTCs while retaining their epithelial characteristics. This phenomenon, also known as the Unjamming Transition (UJT), may happen separately from EMT and contribute to metastasis^36^. As a result, although our simplistic model could infer and predict EMT trajectories at the single-cell resolution, future work needs to consider the complex interactions of tumor cells with the TME to reliably understand the progression of EMT in-vivo.

In conclusion, we introduced COMET, an EMT trajectory inferential and predictive analytic tool. Utilizing COMET, we further resolve the context-specific nature of EMT and argue that the observed pattern of progression is highly dependent on the tumor subtype. Here, we specifically applied COMET to EMT data. However, we anticipate that our general framework is widely applicable for studying phenotypic transitions in other biological processes with intermediate phenotypes.

## 5. Methods

### Flow Cytometry Data

The flow cytometry data was acquired through a Z-Cad dual sensor which was inserted into an MCF10 cell line^22^. This sensor consisted of two components; a green fluorescent protein (GFP) reporter and a red fluorescent protein (RFP) report. While the former was regulated by ZEB, the latter was regulated through E-cad. The fraction of cells in E, H, and M states were found by counting the number of RFP+GFP-, RFP+GFP+, and RFP-GFP+ cells ^22^. Since the RFP-GFP-cells were excluded from counting, the three fractions were normalized for inferring E, H, and M trajectories.

### Gillespie Simulation

Gillespie simulations were performed in Matlab 2021. We simulated the arrival of four exponential random variables with parameters representing the transition rates of the CTMC. At discrete time steps, the system updated with the fastest arriving transition events. There were four transition events from *E* to *H*, *H* to *M*, *M* to *H*, and *H* to *E*. The rates of transitions were drawn from an exponential distribution. We ran simulations until sufficiently enough steps had passed (10000 iterations) and the system had reached steady state. For Figure 3D, the timescale was normalized and trimmed based on the case with the lowest timescale (Treatment withdrawal at day 9). Approximate number of iterations needed to recapitulate Figure3D are: 98, 733, 4000, and 154.

### Gene Set Enrichment Analysis

GSEA was performed using the *fgsea* package^48^ in R 4.2.1. Cells in every cluster were quality controlled and library size normalized. Wilcoxon test was then performed on genes and the resulting rankings were feeded to *fgsea*. All of the reference genes used in this analysis where from the MSigDB database. GSEA was then repeated 10 times for every cluster and the results were overlapped and attached to the results across all clusters in a circular fashion.

## Data Availability

All scripts used in the manuscript are uploaded and available on our Github.

## Acknowledgements

We would like to thank Abhijeet Deshmukh and Sendurai A Mani from UT MD Anderson Cancer Center for providing the raw flow cytometry data used in this study^22^.

JTG was supported by the Cancer Prevention and Research Institute of Texas (RR210080). JTG is a CPRIT Scholars in Cancer Research.

## Software

The Graphical abstract was created with Biorender. All other graphical illustrations were created with Affinity Designer. The mathematical model and simulation were performed using Matlab 2021. The data-driven pipeline, data processing, and data analysis were performed in R 4.2.1.

## Authors’ Contributions

AN performed the research. AN created the computational pipeline AN developed and refined the methodology. AN JTG MKJ analyzed and interpreted the data. AN JTG MKJ wrote the paper. All authors have read and edited the manuscript. JTG conceived of the research. JTG and MKJ designed the research.

## Notes

### Competing Interest Statement

The authors have declared no competing interest.

