## Supplementary Information for "Population Dynamics of EMT Elucidates the Timing and Distribution of Phenotypic Intra-tumoral Heterogeneity"

### 1 Model Development

Our general modeling framework is as follows: We consider a population of fixed size consisting of $N$ total cells divided into 3 sub-populations Epithelial (E), Hybrid (H), and Mesenchymal (M), where H is intermediary to E and M states.

The dynamics of these transitions will be modeled as a continuous-time Markov process, so that the inter-arrival times for each event are exponentially distributed. Transition rates to (resp. from) the $i^{\text{th}}$ state to the next step will be denoted as $\lambda_i$ (resp. $\mu_i$) for $i \in \{E, M\}$ (Figure 1 in main). In general, these transition rates are governed by the cellular environment and as such may vary due to the result of a particular signaling effect $w$, which we assume fixed for a given cell type, as well as the effects of cell spatial location of the EMT phenotype, so that $\lambda_i = \lambda_i(x, y, z; w)$, $\mu_i = \mu_i(x, y, z; w)$. In the results to follow we consider a time inhomogeneous process, which assumes that dynamics are described at some environmental state that is fixed in time. Toward this end, we let the state space consist of the ordered set $\{E, H, M\}$, $E < H < M$. Let $X(t)$ be the random variable denoting the particular state of the population at time $t$. Let $P(t)$ denote the transition matrix of the process, so that $p_{ij}(t) = X(t) = j \mid X(0) = i$ describes the transition probabilities. Let $\pi(t)$ denote the row vector detailing the distribution of states at time $t$.
We may express the evolution of this process via the generator matrix $G$ so that

$$P(t) = e^{tG} = \sum_{j=0}^{\infty} \frac{(tG)^n}{n!}, \tag{1}$$

where the infinitesimal generator matrix, $G$, is given by

$$G = \begin{pmatrix} -\mu_E & \mu_E & 0 \\ \lambda_E & -(\lambda_E + \lambda_M) & \lambda_M \\ 0 & \mu_M & -\mu_M \end{pmatrix}. \tag{2}$$

### 2 General Solution

The Kolmogorov forward equation for this process is

$$\frac{dP}{dt} = P(t)G. \tag{3}$$

Which can be written as:

$$p'_{ij}(t) = \sum_{k \in S} p_{ik}(t) g_{kj} \tag{4}$$

Preprint not peer reviewed

This preprint research paper has not been peer reviewed. Electronic copy available at: https://ssrn.com/abstract=4318055

for all i,j $\in S$

#### 2.1 Equilibrium States

Let $\pi = (\pi_E, \pi_H, \pi_M)$ denote the stationary distribution of the process corresponding to

$$\pi P(t) = \pi; \qquad \pi G = 0. \tag{5}$$

Together with the probability constraint, Eq. 5 corresponding to the following system of equations

$$-\mu_E \pi_E + \lambda_E \pi_H = 0, \tag{6}$$
$$\lambda_M \pi_H - \mu_M \pi_H = 0, \tag{7}$$
$$\pi_E + \pi_H + \pi_M = 1. \tag{8}$$
$$\tag{9}$$

This system admits a unique solution for the stationary distribution, given by

$$\pi = \begin{pmatrix}\pi_E & \pi_H & \pi_M\end{pmatrix} = \begin{pmatrix}\frac{\varphi_E}{1+\varphi_E+\varphi_M} & \frac{1}{1+\varphi_E+\varphi_M} & \frac{\varphi_M}{1+\varphi_E+\varphi_M}\end{pmatrix}, \qquad \text{where } \varphi_i \equiv \lambda_i/\mu_i. \tag{10}$$

### 3 Finding the Transition Probability Matrix

#### 3.1 Eigenvalue Decomposition of the Transition Probability Matrix

The eigenvalues of the transition probability matrix can be found by solving the characteristic equation (Equation 11):

$$\det(G - \eta I) = 0. \tag{11}$$

By definition, $\eta_0 = 0$ is one root that corresponds to $v_0 = \begin{pmatrix}1 & 1 & 1\end{pmatrix}^T$. The remaining are found to be:

$$\eta_{1,2} = \frac{1}{2}\left[-(\lambda_E + \lambda_M + \mu_E + \mu_M) \pm \sqrt{(\lambda_E + \lambda_M + \mu_E - \mu_M)^2 - 4\lambda_M(\mu_E - \mu_M)}\right]. \tag{12}$$

From Equation 12, it is clear that $\eta_1, \eta_2 < 0$, and each can be used to solve for a corresponding eigenvector.



Preprint not peer reviewed

This preprint research paper has not been peer reviewed. Electronic copy available at: https://ssrn.com/abstract=4318055

#### 3.2 Generator eigenvectors

The corresponding eigenvectors are then found to be:

$$v_0 = \begin{bmatrix} 1 & 1 & 1 \end{bmatrix},$$

$$v_1 = \begin{bmatrix} \frac{2\mu_E\lambda_E}{\mu_M(-\mu_E-\lambda_E+\lambda_M+\mu_M+\sqrt{(\mu_E+\lambda_E+\lambda_M+\mu_M)^2-4(\lambda_E\mu_M+\mu_E(\mu_M+\lambda_M))})} \\ -\frac{\lambda_M(-\mu_E+\lambda_E+\lambda_M+\mu_M+\sqrt{(\mu_E+\lambda_E+\lambda_M+\mu_M)^2-4(\lambda_E\mu_M+\mu_E(\mu_M+\lambda_M))})}{\mu_M(-\mu_E-\lambda_E+\lambda_M+\mu_M+\sqrt{(\mu_E+\lambda_E+\lambda_M+\mu_M)^2-4(\lambda_E\mu_M+\mu_E(\mu_M+\lambda_M))})} \\ 1 \end{bmatrix}, \tag{13}$$

$$v_2 = \begin{bmatrix} -\frac{2\mu_E\lambda_E}{\mu_M(\mu_E+\lambda_E-\lambda_M-\mu_M+\sqrt{(\mu_E+\lambda_E+\lambda_M+\mu_M)^2-4(\lambda_E\mu_M+\mu_E(\mu_M+\lambda_M))})} \\ \frac{\lambda_M(-\mu_E+\lambda_E+\lambda_M+\mu_M+\sqrt{(\mu_E+\lambda_E+\lambda_M+\mu_M)^2-4(\lambda_E\mu_M+\mu_E(\mu_M+\lambda_M))})}{\mu_M(\mu_E+\lambda_E-\lambda_M-\mu_M+\sqrt{(\mu_E+\lambda_E+\lambda_M+\mu_M)^2-4(\lambda_E\mu_M+\mu_E(\mu_M+\lambda_M))})} \\ 1 \end{bmatrix},$$

Then we can factorize the generator matrix as

$$G = Q\Lambda Q^{-1}, \tag{14}$$

where we can find matrix Q with $\phi = \sqrt{(\mu_E + \lambda_E + \lambda_M + \mu_M)^2 - 4(\mu_E(\lambda_M + \mu_M) + \lambda_E\mu_M)}$

$$Q = \begin{pmatrix} 1 & \frac{\mu_E(\phi+\mu_E+\lambda_E-\lambda_M-\mu_M)}{2\lambda_E\mu_M} & \frac{\mu_E(-\phi+\mu_E+\lambda_E-\lambda_M-\mu_M)}{2\lambda_E\mu_M} \\ 1 & -\frac{\phi+\mu_E+\lambda_E+\lambda_M-\mu_M}{2\mu_M} & -\frac{-\phi+\mu_E+\lambda_E+\lambda_M-\mu_M}{2\mu_M} \\ 1 & 1 & 1 \end{pmatrix} \tag{15}$$

and $Q^{-1}$ equal to

$$\begin{pmatrix} \frac{\lambda_E\mu_M}{\mu_E(\lambda_M+\mu_M)+\lambda_E\mu_M} & \frac{\mu_E\mu_M}{\mu_E(\lambda_M+\mu_M)+\lambda_E\mu_M} & \frac{\mu_E\lambda_M}{\mu_E(\lambda_M+\mu_M)+\lambda_E\mu_M} \\ \frac{\lambda_E\mu_M(-\phi+\mu_E+\lambda_E+\lambda_M+\mu_M)}{2(\mu_E(\lambda_M+\mu_M)+\lambda_E\mu_M)\phi} & -\frac{\mu_M(-\mu_E^2+\mu_E\phi-\mu_E\lambda_E+\mu_E\lambda_M+\mu_E\mu_M+2\lambda_E\mu_M)}{2(\mu_E(\lambda_M+\mu_M)+\lambda_E\mu_M)\phi} & \frac{\mu_M(-\mu_E^2+\mu_E\phi+\lambda_E\phi-2\mu_E\lambda_E+\mu_E\lambda_M+\mu_E\mu_M-\lambda_E^2-\lambda_E\lambda_M+\lambda_E\mu_M)}{2(\mu_E(\lambda_M+\mu_M)+\lambda_E\mu_M)\phi} \\ -\frac{\lambda_E\mu_M(\phi+\mu_E+\lambda_E+\lambda_M+\mu_M)}{2(\mu_E(\lambda_M+\mu_M)+\lambda_E\mu_M)\phi} & -\frac{\mu_M(\mu_E^2+\mu_E\phi+\mu_E\lambda_E-\mu_E\lambda_M-\mu_E\mu_M-2\lambda_E\mu_M)}{2(\mu_E(\lambda_M+\mu_M)+\lambda_E\mu_M)\phi} & \frac{\mu_M(\mu_E^2+\mu_E\phi+\lambda_E\phi+2\mu_E\lambda_E-\mu_E\lambda_M-\mu_E\mu_M+\lambda_E^2+\lambda_E\lambda_M-\lambda_E\mu_M)}{2(\mu_E(\lambda_M+\mu_M)+\lambda_E\mu_M)\phi} \end{pmatrix} \tag{16}$$

and

$$\Lambda = \begin{pmatrix} 0 & 0 & 0 \\ 0 & \frac{1}{2}(-\phi-\mu_E-\lambda_E-\lambda_M-\mu_M) & 0 \\ 0 & 0 & \frac{1}{2}(\phi-\mu_E-\lambda_E-\lambda_M-\mu_M) \end{pmatrix} \tag{17}$$

We can then consider $\gamma_1 = \frac{1}{2}(-\phi - \mu_E - \lambda_E - \lambda_M - \mu_M)$, $\gamma_2 = \frac{1}{2}(\phi - \mu_E - \lambda_E - \lambda_M - \mu_M)$ and find

$$e^{t\Lambda} = \begin{pmatrix} 1 & 0 & 0 \\ 0 & e^{\gamma_1} & 0 \\ 0 & 0 & e^{\gamma_2} \end{pmatrix} \tag{18}$$



Preprint not peer reviewed

This preprint research paper has not been peer reviewed. Electronic copy available at: https://ssrn.com/abstract=4318055

Where $$\mathrm{e}^{t\lambda}Q^{-1} = \begin{pmatrix} \frac{\lambda_E\mu_M}{\mu_E(\lambda_M+\mu_M)+\lambda_E\mu_M} & \frac{\mu_E\mu_M}{\mu_E(\lambda_M+\mu_M)+\lambda_E\mu_M} & \frac{\mu_E\lambda_M}{\mu_E(\lambda_M+\mu_M)+\lambda_E\mu_M} \\ e^{\gamma_1}\frac{\lambda_E\mu_M(-\phi+\mu_E+\lambda_E+\lambda_M+\mu_M)}{2(\mu_E(\lambda_M+\mu_M)+\lambda_E\mu_M)\phi} & e^{\gamma_1}(-\frac{\mu_M(-\mu_E^2+\mu_E\phi-\mu_E\lambda_E+\mu_E\lambda_M+\mu_E\mu_M+2\lambda_E\mu_M)}{2(\mu_E(\lambda_M+\mu_M)+\lambda_E\mu_M)\phi}) & e^{\gamma_1}\frac{\mu_M(-\mu_E^2+\mu_E\phi+\lambda_E\phi-2\mu_E\lambda_E+\mu_E\lambda_M+\mu_E\mu_M-\lambda_E^2-\lambda_E\lambda_M+\lambda_E\mu_M)}{2(\mu_E(\lambda_M+\mu_M)+\lambda_E\mu_M)\phi} \\ e^{\gamma_2}(-\frac{\lambda_E\mu_M(\phi+\mu_E+\lambda_E+\lambda_M+\mu_M)}{2(\mu_E(\lambda_M+\mu_M)+\lambda_E\mu_M)\phi}) & e^{\gamma_2}(-\frac{\mu_M(\mu_E^2+\mu_E\phi+\mu_E\lambda_E-\mu_E\lambda_M-\mu_E\mu_M-2\lambda_E\mu_M)}{2(\mu_E(\lambda_M+\mu_M)+\lambda_E\mu_M)\phi}) & e^{\gamma_2}(\frac{\mu_M(\mu_E^2+\mu_E\phi+\lambda_E\phi+2\mu_E\lambda_E-\mu_E\lambda_M-\mu_E\mu_M+\lambda_E^2+\lambda_E\lambda_M-\lambda_E\mu_M)}{2(\mu_E(\lambda_M+\mu_M)+\lambda_E\mu_M)\phi}) \end{pmatrix} \quad (19)$$

or find

$$Qe^{t\Lambda} = \begin{pmatrix} 1 & \frac{\mu_E(\phi+\mu_E+\lambda_E-\lambda_M-\mu_M)}{2\lambda_E\mu_M}e^{\frac{1}{2}t(-\phi-\mu_E-\lambda_E-\lambda_M-\mu_M)} & \frac{\mu_E(-\phi+\mu_E+\lambda_E-\lambda_M-\mu_M)}{2\lambda_E\mu_M}e^{\frac{1}{2}t(\phi-\mu_E-\lambda_E-\lambda_M-\mu_M)} \\ 1 & -\frac{\phi+\mu_E+\lambda_E+\lambda_M-\mu_M}{2\mu_M}e^{\frac{1}{2}t(-\phi-\mu_E-\lambda_E-\lambda_M-\mu_M)} & -\frac{-\phi+\mu_E+\lambda_E+\lambda_M-\mu_M}{2\mu_M}e^{\frac{1}{2}t(\phi-\mu_E-\lambda_E-\lambda_M-\mu_M)} \\ 1 & e^{\frac{1}{2}t(-\phi-\mu_E-\lambda_E-\lambda_M-\mu_M)} & e^{\frac{1}{2}t(\phi-\mu_E-\lambda_E-\lambda_M-\mu_M)} \end{pmatrix} \quad (20)$$

The transition probability matrix P with elements as below

$$P = \begin{pmatrix} P_{1,1} & P_{1,2} & P_{1,3} \\ P_{2,1} & P_{2,2} & P_{2,3} \\ P_{3,1} & P_{3,2} & P_{3,3} \end{pmatrix} \quad (21)$$

would then be equivalent to:

$$P_{1,1} = -\frac{\phi}{\mu_M(-\frac{\mu_E\lambda_M\phi}{\lambda_E\mu_M^2}-\frac{\phi}{\mu_M}-\frac{\mu_E\phi}{\lambda_E\mu_M})} + (e^{\gamma_1 * t}(\mu_E^2+\mu_E\lambda_E-\mu_E\lambda_M-\mu_E\mu_M+\mu_E\phi)$$
$$(-\frac{1}{2}-\frac{\mu_E}{2\mu_M}-\frac{\lambda_E}{2\mu_M}-\frac{\lambda_M}{2\mu_M}+\frac{\phi}{2\mu_M}))/(2\lambda_E\mu_M(-\frac{\mu_E\lambda_M\phi}{\lambda_E\mu_M^2}-\frac{\phi}{\mu_M}-\frac{\mu_E\phi}{\lambda_E\mu_M}))+$$
$$(e^{\gamma_2 * t}(\mu_E^2+\mu_E\lambda_E-\mu_E\lambda_M-\mu_E\mu_M-\mu_E\phi)(\frac{1}{2}+\frac{\mu_E}{2\mu_M}+\frac{\lambda_E}{2\mu_M}+\frac{\lambda_M}{2\mu_M}+\frac{\phi}{2\mu_M}))/(2\lambda_E\mu_M(-\frac{\mu_E\lambda_M\phi}{\lambda_E\mu_M^2}-\frac{\phi}{\mu_M}-\frac{\mu_E\phi}{\lambda_E\mu_M}))$$

$$P_{1,2} = -\frac{\mu_E\phi}{\lambda_E\mu_M(-\frac{\mu_E\lambda_M\phi}{\lambda_E\mu_M^2}-\frac{\phi}{\mu_M}-\frac{\mu_E\phi}{\lambda_E\mu_M})} + (e^{\gamma_2 * t}(\mu_E^2+\mu_E\lambda_E-\mu_E\lambda_M-\mu_E\mu_M-\mu_E\phi)$$
$$(-1-\frac{\mu_E}{2\lambda_E}+\frac{\mu_E}{2\mu_M}+\frac{\mu_E^2}{2\lambda_E\mu_M}-\frac{\mu_E\lambda_M}{2\lambda_E\mu_M}+\frac{\mu_E\phi}{2\lambda_E\mu_M}))/(2\lambda_E\mu_M(-\frac{\mu_E\lambda_M\phi}{\lambda_E\mu_M^2}-\frac{\phi}{\mu_M}-\frac{\mu_E\phi}{\lambda_E\mu_M}))+$$
$$(e^{\gamma_1 * t}(\mu_E^2+\mu_E\lambda_E-\mu_E\lambda_M-\mu_E\mu_M+\mu_E\phi)(1+\frac{\mu_E}{2\lambda_E}-\frac{\mu_E}{2\mu_M}-\frac{\mu_E^2}{2\lambda_E\mu_M}+$$
$$\frac{\mu_E\lambda_M}{2\lambda_E\mu_M}+\frac{\mu_E\phi}{2\lambda_E\mu_M}))/(2\lambda_E\mu_M(-\frac{\mu_E\lambda_M\phi}{\lambda_E\mu_M^2}-\frac{\phi}{\mu_M}-\frac{\mu_E\phi}{\lambda_E\mu_M}))$$

$$P_{1,3} = -\frac{\mu_E\lambda_M\phi}{\lambda_E\mu_M^2(-\frac{\mu_E\lambda_M\phi}{\lambda_E\mu_M^2}-\frac{\phi}{\mu_M}-\frac{\mu_E\phi}{\lambda_E\mu_M})} + (e^{\gamma_1 * t}(\mu_E^2+\mu_E\lambda_E-\mu_E\lambda_M-\mu_E\mu_M+\mu_E\phi)$$
$$(-\frac{1}{2}-\frac{\mu_E}{2\lambda_E}+\frac{\mu_E}{\mu_M}+\frac{\mu_E^2}{2\lambda_E\mu_M}+\frac{\lambda_E}{2\mu_M}+\frac{\lambda_M}{2\mu_M}-\frac{\mu_E\lambda_M}{2\lambda_E\mu_M}-\frac{\phi}{2\mu_M}-\frac{\mu_E\phi}{2\lambda_E\mu_M}))/(2\lambda_E\mu_M(-\frac{\mu_E\lambda_M\phi}{\lambda_E\mu_M^2}-\frac{\phi}{\mu_M}-\frac{\mu_E\phi}{\lambda_E\mu_M}))+$$
$$(e^{\gamma_2 * t}(\mu_E^2+\mu_E\lambda_E-\mu_E\lambda_M-\mu_E\mu_M-\mu_E\phi)(\frac{1}{2}+\frac{\mu_E}{2\lambda_E}-\frac{\mu_E}{\mu_M}-\frac{\mu_E^2}{2\lambda_E\mu_M}-$$
$$\frac{\lambda_E}{2\mu_M}-\frac{\lambda_M}{2\mu_M}+\frac{\mu_E\lambda_M}{2\lambda_E\mu_M}-\frac{\phi}{2\mu_M}-\frac{\mu_E\phi}{2\lambda_E\mu_M}))/(2\lambda_E\mu_M(-\frac{\mu_E\lambda_M\phi}{\lambda_E\mu_M^2}-\frac{\phi}{\mu_M}-\frac{\mu_E\phi}{\lambda_E\mu_M}))$$

Preprint not peer reviewed



This preprint research paper has not been peer reviewed. Electronic copy available at: https://ssrn.com/abstract=4318055

$$P_{2,1} = -\frac{\phi}{\mu_M(-\frac{\mu_E\lambda_M\phi}{\lambda_E\mu_M^2} - \frac{\phi}{\mu_M} - \frac{\mu_E\phi}{\lambda_E\mu_M})} + (e^{-\frac{1}{2}(\mu_E+\lambda_E+\lambda_M+\mu_M+\phi)t}(-\mu_E - \lambda_E - \lambda_M + \mu_M - \phi)$$
$$(-\frac{1}{2} - \frac{\mu_E}{2\mu_M} - \frac{\lambda_E}{2\mu_M} - \frac{\lambda_M}{2\mu_M} + \frac{\phi}{2\mu_M}))/(2\mu_M(-\frac{\mu_E\lambda_M\phi}{\lambda_E\mu_M^2} - \frac{\phi}{\mu_M} - \frac{\mu_E\phi}{\lambda_E\mu_M}))+$$
$$(e^{\gamma_2 * t}(-\mu_E-\lambda_E-\lambda_M+\mu_M+\phi)(\frac{1}{2}+\frac{\mu_E}{2\mu_M}+\frac{\lambda_E}{2\mu_M}+\frac{\lambda_M}{2\mu_M}+\frac{\phi}{2\mu_M}))/(2\mu_M(-\frac{\mu_E\lambda_M\phi}{\lambda_E\mu_M^2}-\frac{\phi}{\mu_M}-\frac{\mu_E\phi}{\lambda_E\mu_M}))$$

$$P_{2,2} = -\frac{\mu_E\phi}{\lambda_E\mu_M(-\frac{\mu_E\lambda_M\phi}{\lambda_E\mu_M^2} - \frac{\phi}{\mu_M} - \frac{\mu_E\phi}{\lambda_E\mu_M})} + (e^{\gamma_2 * t}(-\mu_E - \lambda_E - \lambda_M + \mu_M + \phi)$$
$$(-1 - \frac{\mu_E}{2\lambda_E} + \frac{\mu_E}{2\mu_M} + \frac{\mu_E^2}{2\lambda_E\mu_M} - \frac{\mu_E\lambda_M}{2\lambda_E\mu_M} + \frac{\mu_E\phi}{2\lambda_E\mu_M}))/(2\mu_M(-\frac{\mu_E\lambda_M\phi}{\lambda_E\mu_M^2} - \frac{\phi}{\mu_M} - \frac{\mu_E\phi}{\lambda_E\mu_M}))+$$
$$(e^{\gamma_1 * t}(-\mu_E - \lambda_E - \lambda_M + \mu_M - \phi)(1 + \frac{\mu_E}{2\lambda_E} - \frac{\mu_E}{2\mu_M} - \frac{\mu_E^2}{2\lambda_E\mu_M}+$$
$$\frac{\mu_E\lambda_M}{2\lambda_E\mu_M} + \frac{\mu_E\phi}{2\lambda_E\mu_M}))/(2\mu_M(-\frac{\mu_E\lambda_M\phi}{\lambda_E\mu_M^2} - \frac{\phi}{\mu_M} - \frac{\mu_E\phi}{\lambda_E\mu_M}))$$

$$P_{2,3} = -\frac{\mu_E\lambda_M\phi}{\lambda_E\mu_M^2(-\frac{\mu_E\lambda_M\phi}{\lambda_E\mu_M^2} - \frac{\phi}{\mu_M} - \frac{\mu_E\phi}{\lambda_E\mu_M})} + (e^{\gamma_1 * t}(-\mu_E - \lambda_E - \lambda_M + \mu_M - \phi)$$
$$(-\frac{1}{2}-\frac{\mu_E}{2\lambda_E}+\frac{\mu_E}{\mu_M}+\frac{\mu_E^2}{2\lambda_E\mu_M}+\frac{\lambda_E}{2\mu_M}+\frac{\lambda_M}{2\mu_M}-\frac{\mu_E\lambda_M}{2\lambda_E\mu_M}-\frac{\phi}{2\mu_M}-\frac{\mu_E\phi}{2\lambda_E\mu_M}))/(2\mu_M(-\frac{\mu_E\lambda_M\phi}{\lambda_E\mu_M^2}-\frac{\phi}{\mu_M}-\frac{\mu_E\phi}{\lambda_E\mu_M}))+$$
$$(e^{\gamma_2 * t}(-\mu_E - \lambda_E - \lambda_M + \mu_M + \phi)(\frac{1}{2} + \frac{\mu_E}{2\lambda_E} - \frac{\mu_E}{\mu_M} - \frac{\mu_E^2}{2\lambda_E\mu_M} -$$
$$\frac{\lambda_E}{2\mu_M} - \frac{\lambda_M}{2\mu_M} + \frac{\mu_E\lambda_M}{2\lambda_E\mu_M} - \frac{\phi}{2\mu_M} - \frac{\mu_E\phi}{2\lambda_E\mu_M}))/(2\mu_M(-\frac{\mu_E\lambda_M\phi}{\lambda_E\mu_M^2} - \frac{\phi}{\mu_M} - \frac{\mu_E\phi}{\lambda_E\mu_M}))$$

$$P_{3,1} = -\frac{\phi}{\mu_M(-\frac{\mu_E\lambda_M\phi}{\lambda_E\mu_M^2} - \frac{\phi}{\mu_M} - \frac{\mu_E\phi}{\lambda_E\mu_M})} + (e^{\gamma_1 * t}(-\frac{1}{2} - \frac{\mu_E}{2\mu_M} - \frac{\lambda_E}{2\mu_M} - \frac{\lambda_M}{2\mu_M}+$$
$$\frac{\phi}{2\mu_M}))/(-\frac{\mu_E\lambda_M\phi}{\lambda_E\mu_M^2} - \frac{\phi}{\mu_M} - \frac{\mu_E\phi}{\lambda_E\mu_M}) + \frac{e^{\gamma_2 * t}(\frac{1}{2} + \frac{\mu_E}{2\mu_M} + \frac{\lambda_E}{2\mu_M} + \frac{\lambda_M}{2\mu_M} + \frac{\phi}{2\mu_M})}{-\frac{\mu_E\lambda_M\phi}{\lambda_E\mu_M^2} - \frac{\phi}{\mu_M} - \frac{\mu_E\phi}{\lambda_E\mu_M}}$$

$$P_{3,2} = -\frac{\mu_E\phi}{\lambda_E\mu_M(-\frac{\mu_E\lambda_M\phi}{\lambda_E\mu_M^2} - \frac{\phi}{\mu_M} - \frac{\mu_E\phi}{\lambda_E\mu_M})} + (e^{\gamma_2 * t}(-1 - \frac{\mu_E}{2\lambda_E} + \frac{\mu_E}{2\mu_M}\frac{\mu_E^2}{2\lambda_E\mu_M}-$$
$$\frac{\mu_E\lambda_M}{2\lambda_E\mu_M}+\frac{\mu_E\phi}{2\lambda_E\mu_M}))/(-\frac{\mu_E\lambda_M\phi}{\lambda_E\mu_M^2}-\frac{\phi}{\mu_M}-\frac{\mu_E\phi}{\lambda_E\mu_M})+(e^{\gamma_1 * t}(1+\frac{\mu_E}{2\lambda_E}-\frac{\mu_E}{2\mu_M}-\frac{\mu_E^2}{2\lambda_E\mu_M}+\frac{\mu_E\lambda_M}{2\lambda_E\mu_M}+\frac{\mu_E\phi}{2\lambda_E\mu_M}))/$$
$$(-\frac{\mu_E\lambda_M\phi}{\lambda_E\mu_M^2} - \frac{\phi}{\mu_M} - \frac{\mu_E\phi}{\lambda_E\mu_M})$$



Preprint not peer reviewed

This preprint research paper has not been peer reviewed. Electronic copy available at: https://ssrn.com/abstract=4318055

$$P_{3,3} = -\frac{\mu_E\lambda_M\phi}{\lambda_E\mu_M^2(-\frac{\mu_E\lambda_M\phi}{\lambda_E\mu_M^2} - \frac{\phi}{\mu_M} - \frac{\mu_E\phi}{\lambda_E\mu_M})} + (e^{\gamma_1 * t}(-\frac{1}{2} - \frac{\mu_E}{2\lambda_E} + \frac{\mu_E}{\mu_M} + \frac{\mu_E^2}{2\lambda_E\mu_M} + \frac{\lambda_E}{2\mu_M} + \frac{\lambda_M}{2\mu_M}$$
$$- \frac{\mu_E\lambda_M}{2\lambda_E\mu_M} - \frac{\phi}{2\mu_M} - \frac{\mu_E\phi}{2\lambda_E\mu_M}))/(-\frac{\mu_E\lambda_M\phi}{\lambda_E\mu_M^2} - \frac{\phi}{\mu_M} - \frac{\mu_E\phi}{\lambda_E\mu_M})+$$
$$(e^{\frac{1}{2}(-\mu_E-\lambda_E-\lambda_M-\mu_M+\phi)t}(\frac{1}{2} + \frac{\mu_E}{2\lambda_E} - \frac{\mu_E}{\mu_M} - \frac{\mu_E^2}{2\lambda_E\mu_M} - \frac{\lambda_E}{2\mu_M} - \frac{\lambda_M}{2\mu_M} + \frac{\mu_E\lambda_M}{2\lambda_E\mu_M} - \frac{\phi}{2\mu_M} - \frac{\mu_E\phi}{2\lambda_E\mu_M}))/$$
$$(-\frac{\mu_E\lambda_M\phi}{\lambda_E\mu_M^2} - \frac{\phi}{\mu_M} - \frac{\mu_E\phi}{\lambda_E\mu_M})$$

Below, we provide the explicit solution in the case where the effects of a phenotypic stability factor are symmetric between E and M phenotypes.

#### 3.3 Symmetric Effects of Phenotypic Stability Factors

Considering that PSFs stabilize the hybrid phenotype in a symmetric fashion, then we may take $\mu_E = \mu_M \equiv \mu$. In this case, the generator becomes

$$G = \begin{pmatrix} -\mu & \mu & 0 \\ \lambda_E & -(\lambda_E + \lambda_M) & \lambda_M \\ 0 & \mu & -\mu \end{pmatrix} \tag{22}$$

and Equation 12 simplifies considerably to

$$\eta_{1,2} = \frac{1}{2}\left[-(\lambda_E + \lambda_M + 2\mu) \pm \sqrt{(\lambda_E + \lambda_M)^2}\right], \tag{23}$$

giving

$$\eta_1 = -\mu; \qquad \eta_2 = -(\lambda_E + \lambda_M + \mu). \tag{24}$$

The corresponding eigenvectors solve

$$\begin{pmatrix} 0 & \mu & 0 \\ \lambda_E & -(\lambda_E + \lambda_M - \mu) & \lambda_M \\ 0 & \mu & 0 \end{pmatrix} v_1 = 0; \qquad \begin{pmatrix} \lambda_E + \lambda_M & \mu & 0 \\ \lambda_E & -\mu & \lambda_M \\ 0 & \mu & \lambda_E + \lambda_M \end{pmatrix} v_2 = 0. \tag{25}$$

Thus, we may take the diagonalization matrix, $Q = \begin{pmatrix} v_0 & v_1 & v_2 \end{pmatrix}$, as

$$Q = \begin{pmatrix} 1 & \lambda_M & -\mu \\ 1 & 0 & \lambda_E + \lambda_M \\ 1 & -\lambda_E & -\mu \end{pmatrix} \tag{26}$$

Note that $\nu \equiv \det Q$ may be calculated by expansion along the first row of $Q$:

$$\nu == 1\begin{vmatrix} 0 & \lambda_E + \lambda_M \\ -\lambda_E & -\mu \end{vmatrix} - \lambda_M \begin{vmatrix} 1 & \lambda_E + \lambda_M \\ 1 & -\mu \end{vmatrix} + (-\mu)\begin{vmatrix} 1 & 0 \\ 1 & -\lambda_E \end{vmatrix} \tag{27}$$
$$= (\lambda_E + \lambda_M)(\lambda_E + \lambda_M + \mu) > 0. \tag{28}$$



Preprint not peer reviewed

This preprint research paper has not been peer reviewed. Electronic copy available at: https://ssrn.com/abstract=4318055

$Q$ is therefore invertible. Its inverse may be calculated as

$$Q^{-1} = \frac{1}{\nu}\begin{pmatrix} \lambda_E(\lambda_E+\lambda_M) & -\mu(\lambda_E-\lambda_M) & -\lambda_M(\lambda_E+\lambda_M) \\ \lambda_E+\lambda_M+\mu & 0 & -(\lambda_E+\lambda_M+\mu) \\ -\lambda_E & -(\lambda_E-\lambda_M) & \lambda_M \end{pmatrix}. \quad (29)$$

$Q$ and $Q^{-1}$ can be used to diagonalize the infinitesimal generator via:

$$G = Q\Lambda Q^{-1}, \quad (30)$$

where

$$\Lambda = \begin{pmatrix} 0 & 0 & 0 \\ 0 & -\mu & 0 \\ 0 & 0 & -(\lambda_E+\lambda_M+\mu) \end{pmatrix}. \quad (31)$$

Clearly,

$$G^n = Q\Lambda^n Q^{-1}. \quad (32)$$

Therefore, we have

$$P(t) = \sum_{j=0}^{\infty} \frac{t^n}{n!} Q\Lambda^n Q^{-1} = Qe^{t\Lambda}Q^{-1}. \quad (33)$$

Putting

$$\begin{aligned} k_1 &= -\eta_1 = \mu \\ k_2 &= -\eta_2 = \lambda_E+\lambda_M+\mu \\ k_3 &= k_2 - k_1 = \lambda_E+\lambda_M, \end{aligned} \quad (34)$$

evaluating Eq. 33, and expressing the result in terms of the stationary distribution from Eq. 10, we may express the exact distribution by

$$P = \begin{pmatrix} \pi_E + e^{-k_1 t}\left[1 - \frac{\lambda_E}{k_3}\left(1-\pi_H e^{-k_3 t}\right)\right] & \pi_H\left(1-e^{-k_2 t}\right) & \pi_M - e^{-k_1 t}\left[\pi_M + \frac{\lambda_M}{k_3}\pi_H\left(1-e^{-k_3 t}\right)\right] \\ \pi_E\left(1-e^{-k_2 t}\right) & \pi_H + e^{-k_2 t}\left(1-\pi_H\right) & \pi_M\left(1-e^{-k_2 t}\right) \\ \pi_E - e^{-k_1 t}\left[\pi_E + \frac{\lambda_E}{k_3}\pi_H\left(1-e^{-k_3 t}\right)\right] & \pi_H\left(1-e^{-k_2 t}\right) & \pi_M + e^{-k_1 t}\left[1 - \frac{\lambda_M}{k_3}\left(1-\pi_H e^{-k_3 t}\right)\right]. \end{pmatrix} \quad (35)$$

### 4 Application to spatial EMT environments

The given dynamics of the process imply that extinction does not occur for any given phenotype, but rather that such cell sizes may be found in very large or small abundances as a function of the relative transition rates. Understanding and predicting their relative



Preprint not peer reviewed

This preprint research paper has not been peer reviewed. Electronic copy available at: https://ssrn.com/abstract=4318055

abundance has direct clinical and experimental relevance to studying cancer, where predicting the distribution of hybrid and mesenchymal cells is useful for predicting cancer aggressiveness and treatment resistance. With this motivation in mind, we consider several applications. We may consider direct exogeneous signals, such as the growth factor TGF$\beta$ present during EMT [1], as well as indirect ones, including other inflammatory cytokines that result in augmented SNAIL concentrations via inflammatory NF$\kappa$B and TNF$\alpha$ signaling at the boundary of a wound or invading cancer population [2]. Toward this end, we consider a 1-dimensional signaling environment (for example, TGF$\beta$ concentration). We will assume that these signals may be represented generally by a sigmoidal function and that their effects impact inversely the epithelial and mesenchymal transition rates. This may generally be represented by the following functional forms:

$$\lambda_E(x) = \lambda_E^+ - \frac{(\lambda_E^+ - \lambda_E^-)x^n}{L + x^n}; \qquad \lambda_M(x) = \lambda_M^- + \frac{(\lambda_M^+ - \lambda_M^-)x^n}{L + x^n}. \tag{36}$$

If we assume $\lambda_E^+ + \lambda_M^- = \lambda_E^- + \lambda_M^+ \equiv C$, then $\lambda_E + \lambda_M = C$. Together with the assumption that the rate of transition into state $H$, $\mu$, is constant and given the fact that the stationary distribution of hybrid cells is $\pi_{H,\infty} = \mu/(\lambda_E + \lambda_M + \mu)$, we can conclude that $\pi_H$ is constant in space (Figure SI.1).

### 5 Dimensionality Analysis of TGF$\beta$ Dose

Considering the molecular weight of TGF$\beta$ is 44kDa, then the dose of TGF$\beta$ is $5 * 10^-9 g/ml * 1 \frac{mol}{44 * 10^3 g} = 0.1136 * 10^{-12} \frac{mol}{ml}$ which is equivalent to $X = 113.6pM$.



Preprint not peer reviewed

This preprint research paper has not been peer reviewed. Electronic copy available at: https://ssrn.com/abstract=4318055

### 6 Supplemental Figures

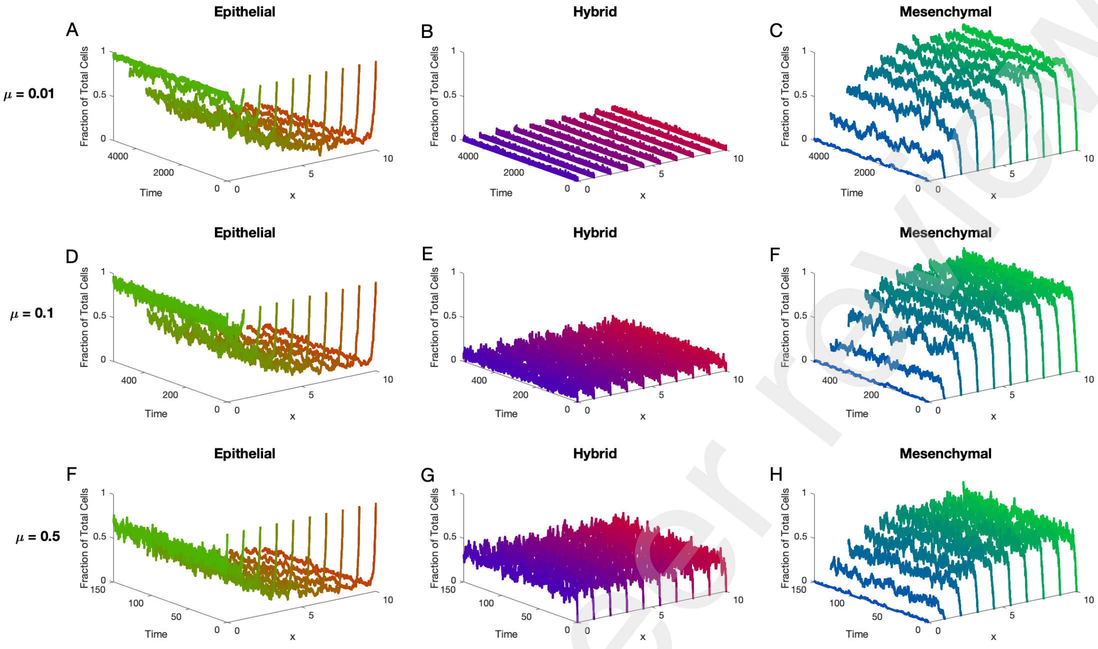


Figure SI.1: **Spatial EMT Environments.** The hybrid state is stabilized in space.



Preprint not peer reviewed

This preprint research paper has not been peer reviewed. Electronic copy available at: https://ssrn.com/abstract=4318055

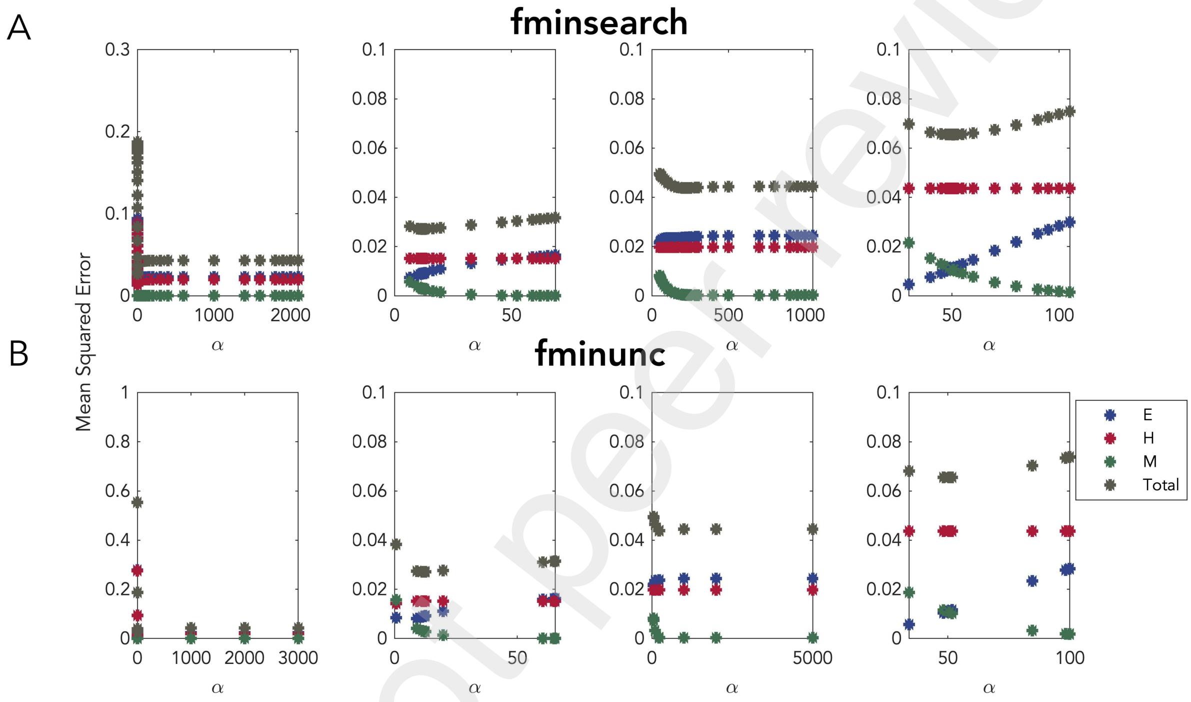


Figure SI.2: **Figure shows the alpha parameter optimized using two Matlab functions; fminsearch and fminunc.** Panel (A) plots show the MSE found using fminsearch while plots of panel (B) show the MSE found using fminunc. These MSEs correspond to Figure 3C in main. Plot shows an overall faster convergence to the optimal value using fminunc.



This preprint research paper has not been peer reviewed. Electronic copy available at: https://ssrn.com/abstract=4318055

Preprint not peer reviewed

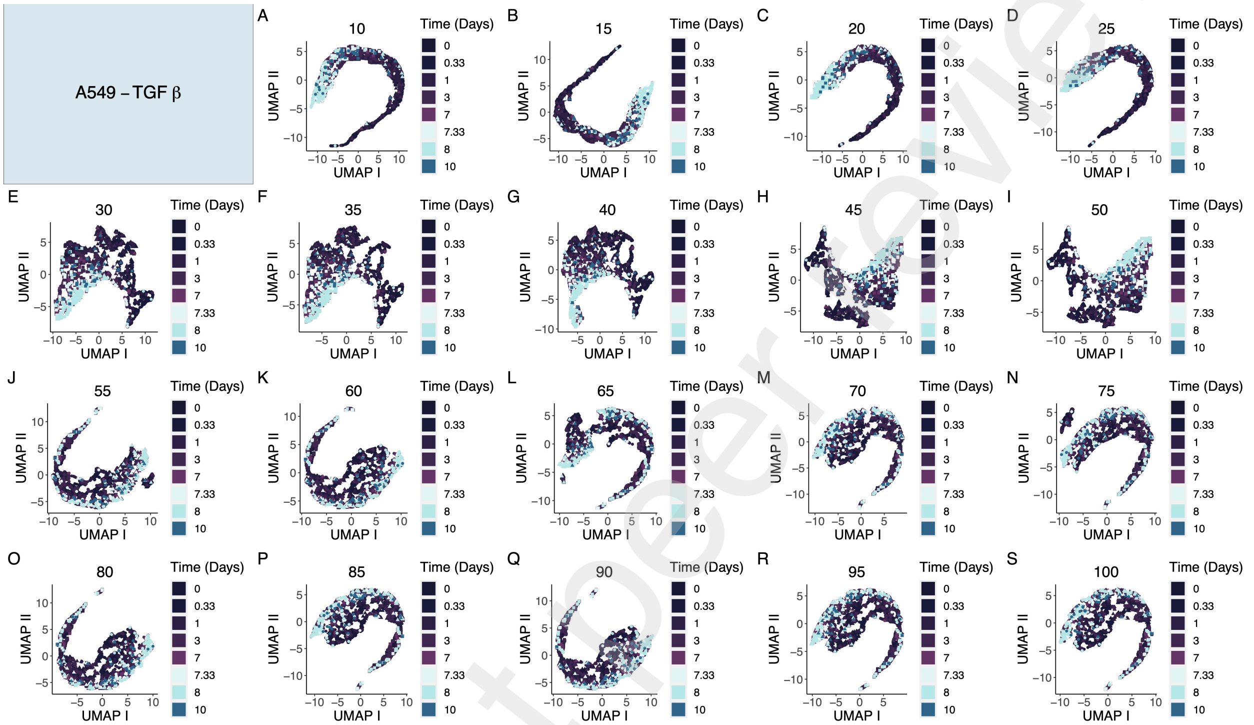


Figure SI.3: **A549 dimension reduction shows the loss of EMT order when the number of included EMT related genes increases.** Figure shows the UMAP plot for one run of the algorithm for different cutoffs of highly variable genes (10-100 in increments of 5). Figure shows a gradual movement from epithelial to mesenchymal phenotype as a function of time during TGF$\beta$ treatment for the lower cutoffs which then trace the same footsteps backwards following treatment withdrawal. This spectrum-like pattern clearly is not captured from data when the number of highly variable genes increases to 50 and above.

Preprint not peer reviewed



This preprint research paper has not been peer reviewed. Electronic copy available at: https://ssrn.com/abstract=4318055

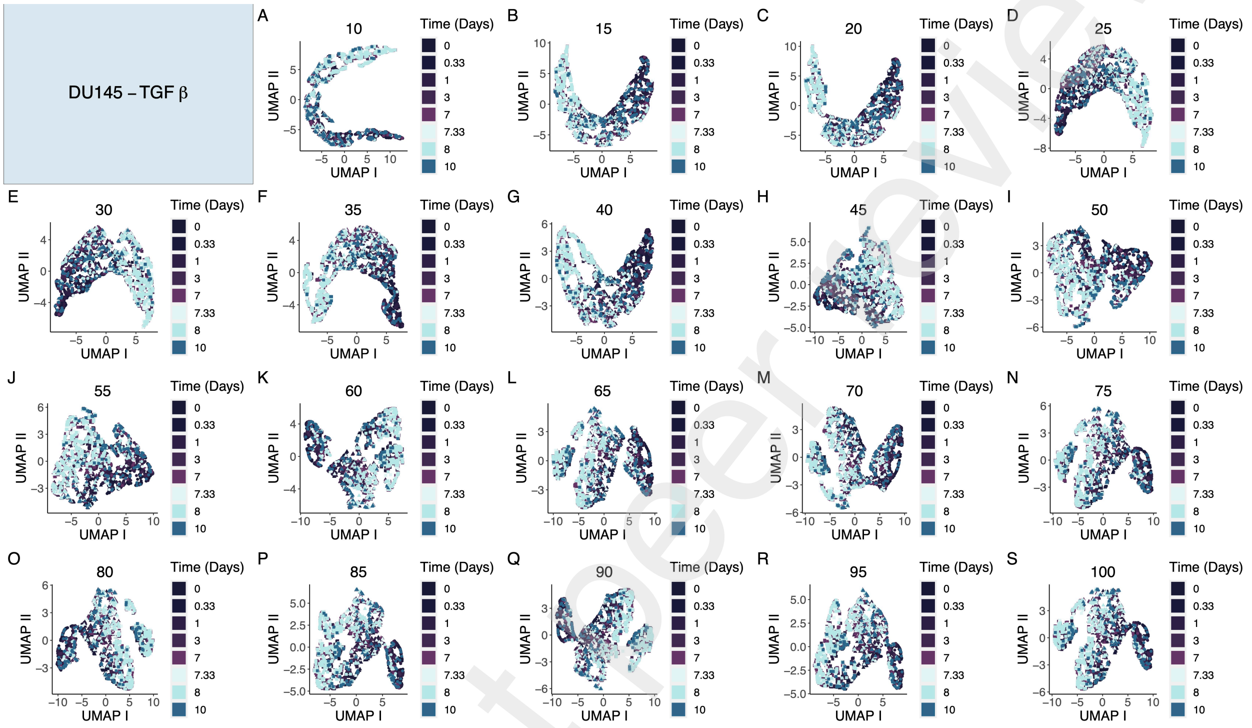


Figure SI.4: **DU145 dimension reduction shows a possible increase in the number of intermediary phenotypes as the number of highly variable genes included is increased.** Figure shows the UMAP plot for one run of the algorithm for different cutoffs of highly variable genes (10-100 in increments of 5). Figure shows a gradual movement from epithelial to mesenchymal phenotype as a function of time during TGF$\beta$ treatment for the lower cutoffs which then trace the same footsteps backwards following treatment withdrawal. This spectrum-like pattern appears to move through more hybrid phenotypes as the number of highly variable genes included in the analysis increases.



This preprint research paper has not been peer reviewed. Electronic copy available at: https://ssrn.com/abstract=4318055

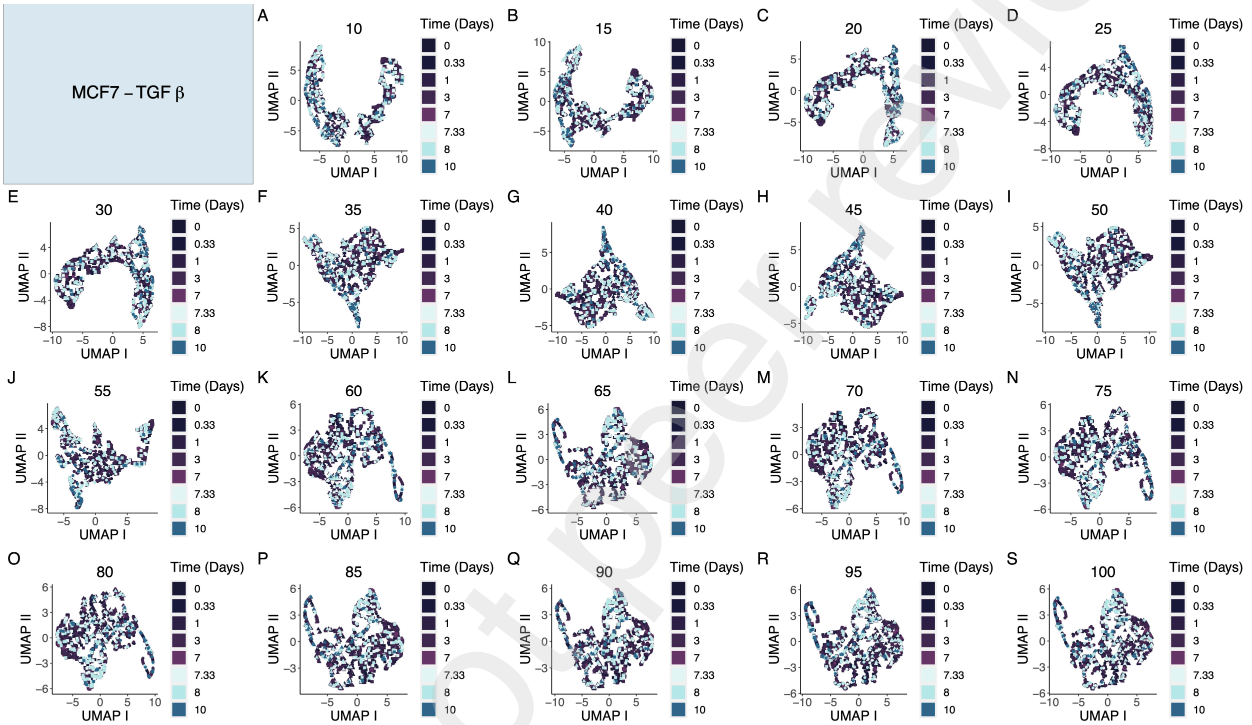

Figure SI.5: **Figure shows significant heterogeneity regardless of cutoff in the UMAP plot.** Figure depicts the UMAP plots for one run of the algorithm for MCF7 treated with TGF$\beta$. Clearly, the pipeline is either detecting another phenomenon from the data, or cells do not go through EMT and subsequent MET.



Preprint not peer reviewed

This preprint research paper has not been peer reviewed. Electronic copy available at: https://ssrn.com/abstract=4318055

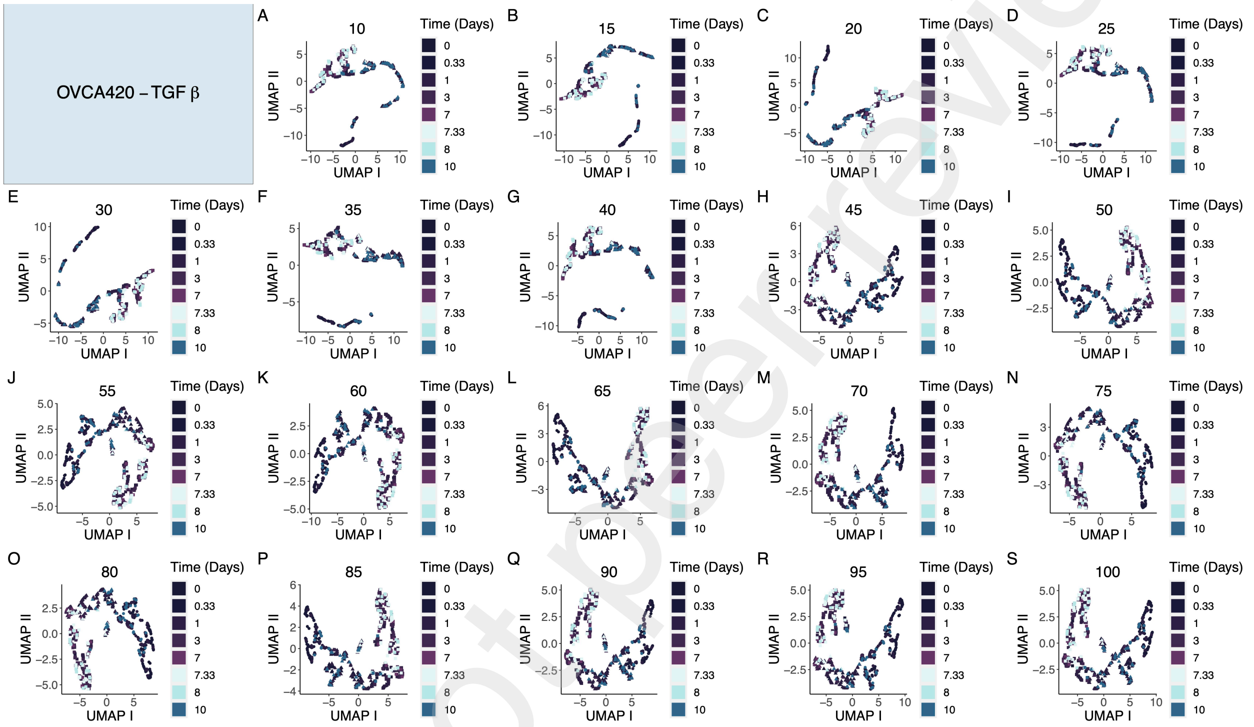


Figure SI.6: **Figure shows the movement of cells from the epithelial cluster to the mesenchymal cluster for OVCA420 treated with TGF$\beta$ when higher number of EMT related genes are included.** Unlike the DU145 case, figure depicts the loss of the spectrum shape of UMAP plot as the number of included genes decreases for lower cutoffs.

Preprint not peer reviewed



This preprint research paper has not been peer reviewed. Electronic copy available at: https://ssrn.com/abstract=4318055

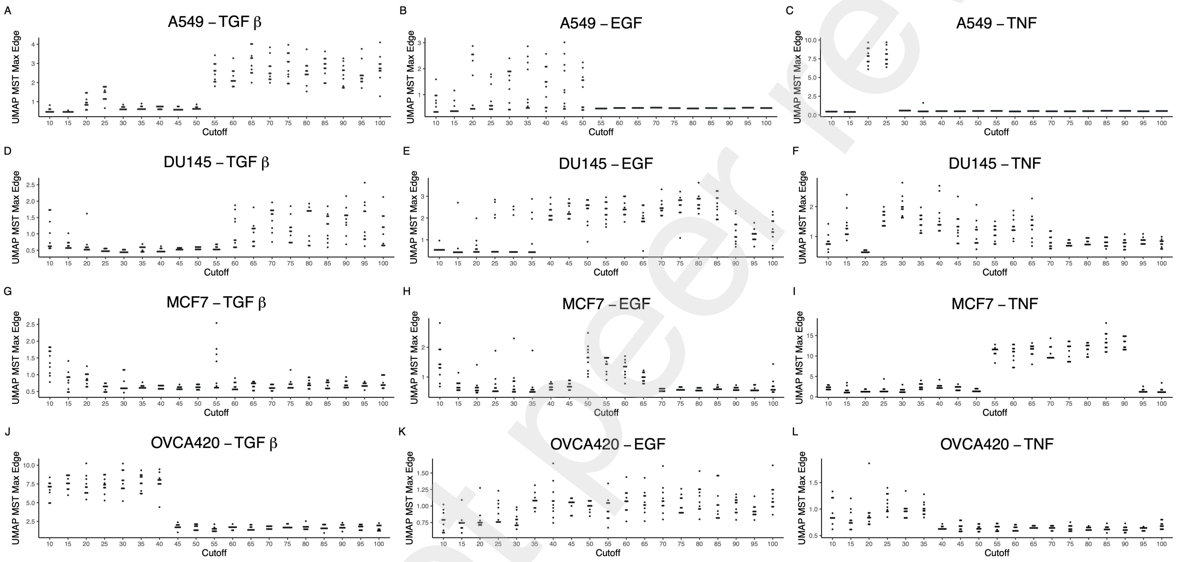


Figure SI.7: **Maximum edge of the minimum spanning tree fitted to data can determine the optimal cutoff.** Plot depicting the maximum edge of the minimum spanning tree.

.

Preprint not peer reviewed



This preprint research paper has not been peer reviewed. Electronic copy available at: https://ssrn.com/abstract=4318055

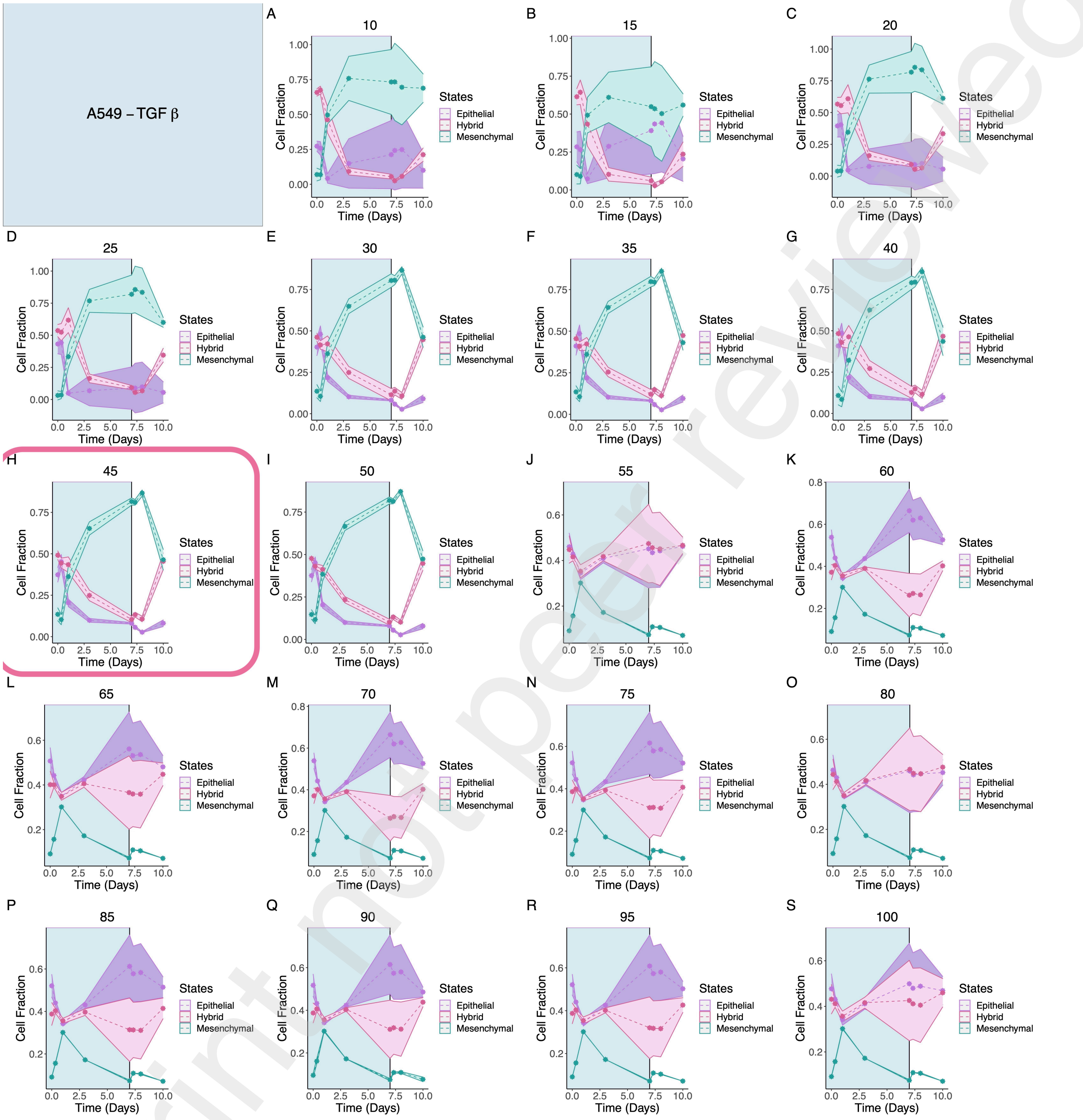


Figure SI.8: **Time-course trajectories of A549 treated with TGF$\beta$.** From the figure, it is clear that the inclusion of only five extra highly variable genes from 50 to 55 can drastically change the trajectories perhaps due to the difference in number of states. The pink box around the plot shows the case with the optimal number of highly variable EMT genes for our three state system based on DTW alignment to flow cytometry data of Jia et al [3].



Preprint not peer reviewed

This preprint research paper has not been peer reviewed. Electronic copy available at: https://ssrn.com/abstract=4318055

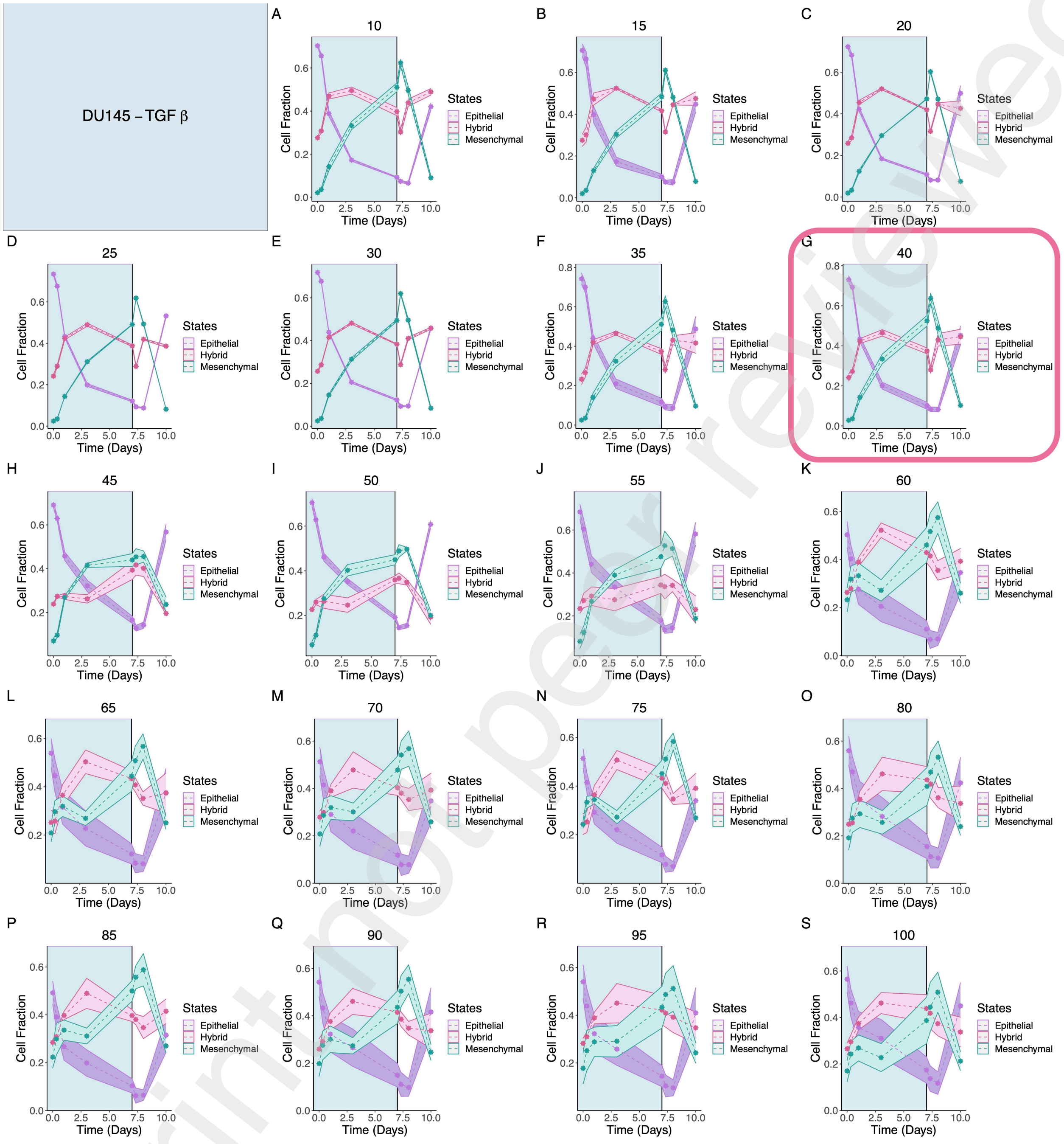


Figure SI.9: **Time-course trajectories of DU145 treated with TGF$\beta$.** Figure shows that collectively the inclusion of more EMT genes result in higher variability and lower resolvability for inferring the three states. The pink box around the plot shows the case with the optimal number of highly variable EMT genes for our three state system based on DTW alignment to flow cytometry data of Jia et al [3].

Preprint not peer reviewed



This preprint research paper has not been peer reviewed. Electronic copy available at: https://ssrn.com/abstract=4318055

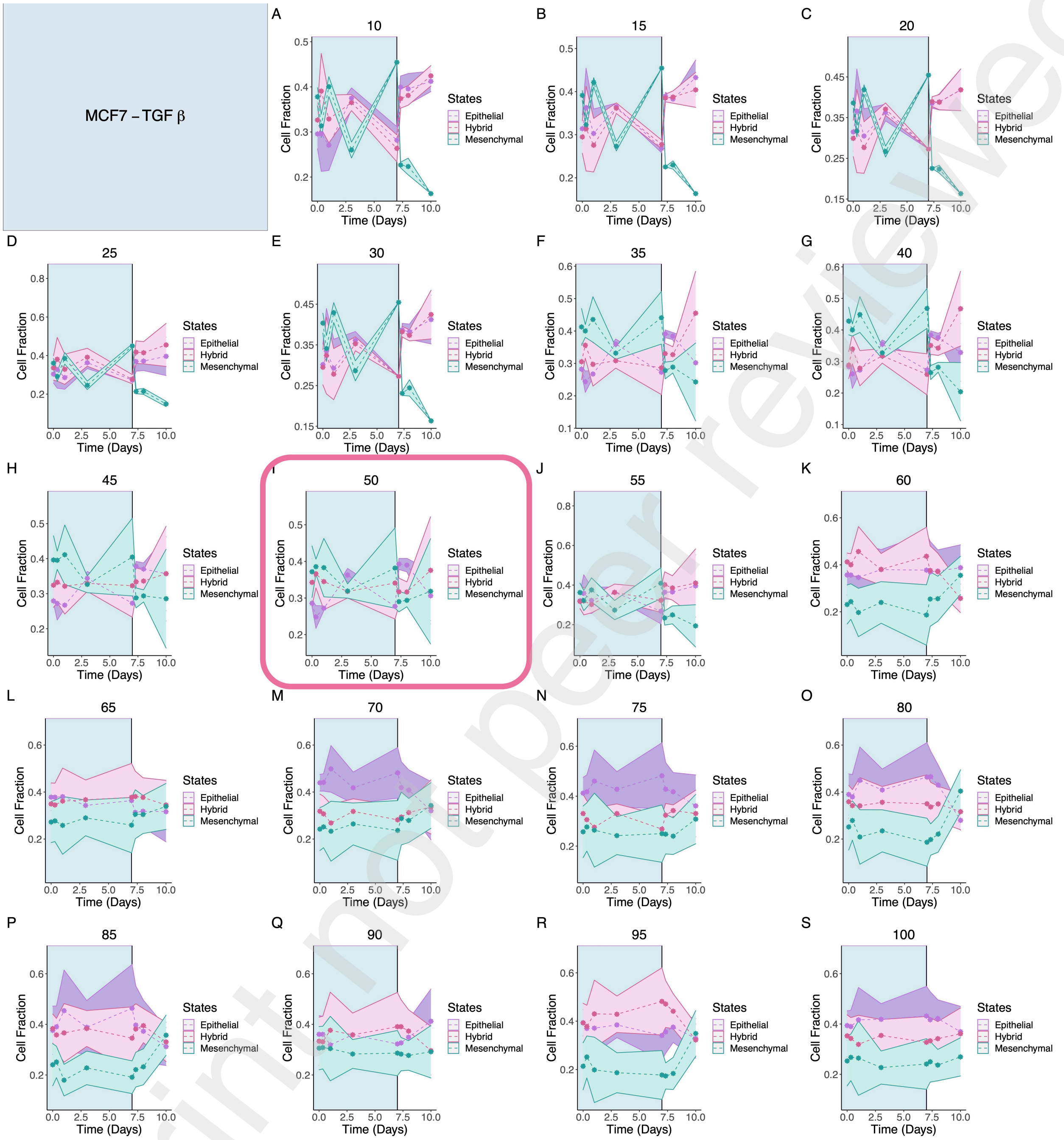


Figure SI.10: **Time-course trajectories of MCF7 treated with TGF$\beta$.** This figure shows that our pipeline likely failed to infer the correct trajectories from the single-cell RNA sequencing pipeline. The pink box around the plot shows the case with the optimal number of highly variable EMT genes for our three state system based on DTW alignment to flow cytometry data of Jia et al [3].



Preprint not peer reviewed

This preprint research paper has not been peer reviewed. Electronic copy available at: https://ssrn.com/abstract=4318055

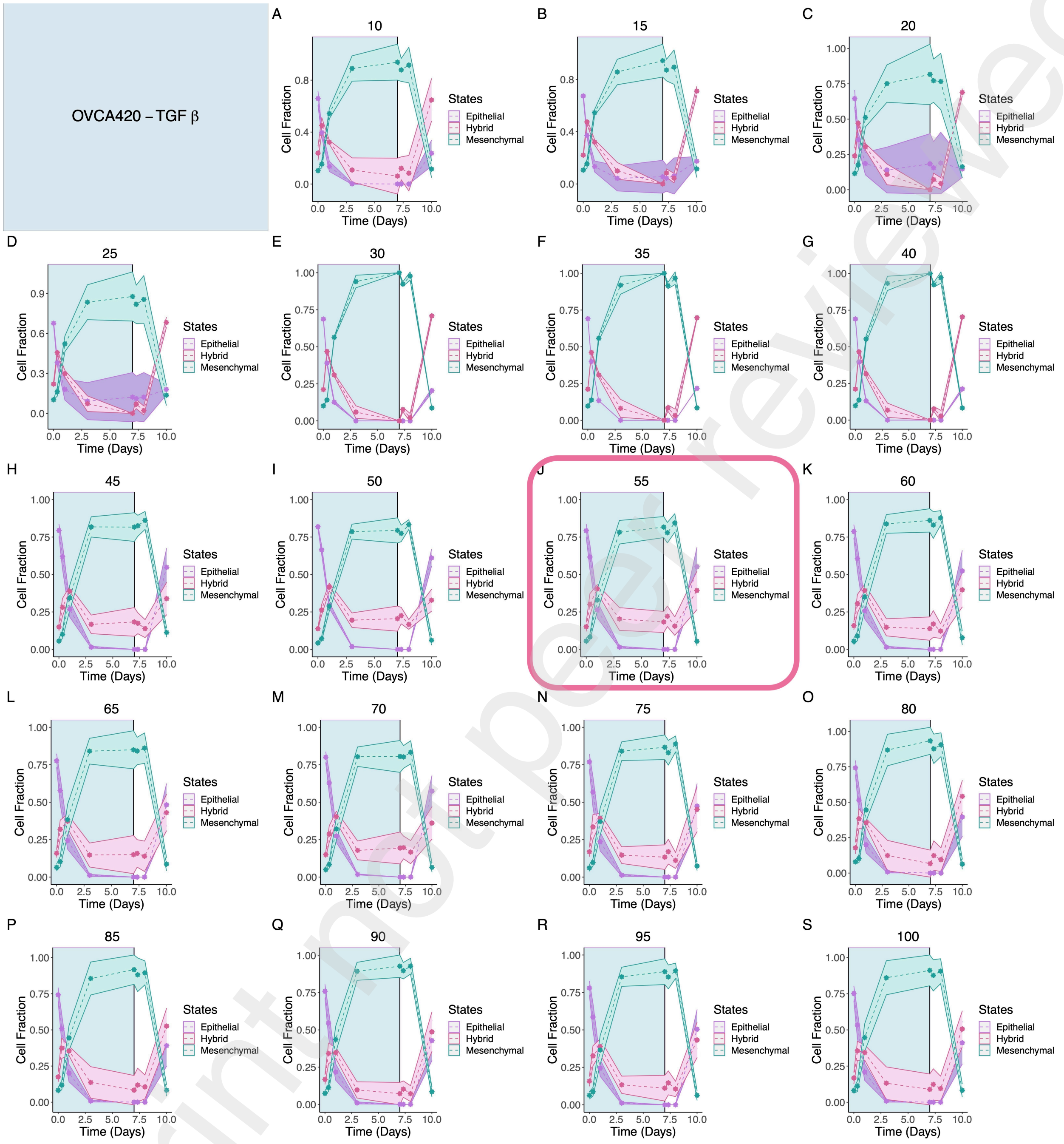


Figure SI.11: **Time-course trajectories of OVCA420 treated with TGF$\beta$.** Figure illustrates that the OVCA420 cell line is mostly epithelial and quickly transitions into a mesenchymal state, residing in the hybrid state only for a short time. The pink box around the plot shows the case with the optimal number of highly variable EMT genes for our three state system based on DTW alignment to flow cytometry data of Jia et al [3].



Preprint not peer reviewed

This preprint research paper has not been peer reviewed. Electronic copy available at: https://ssrn.com/abstract=4318055

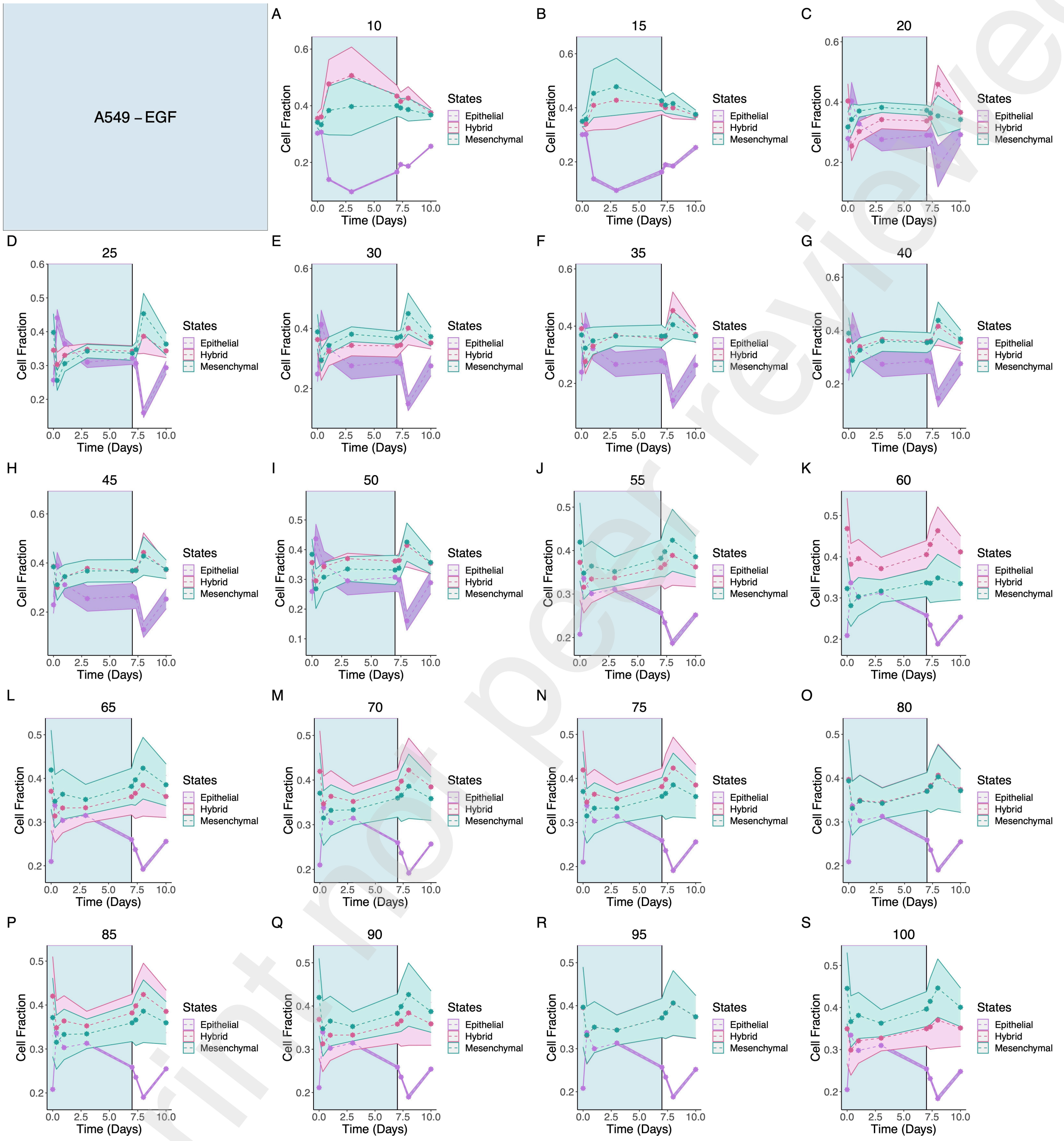


Figure SI.12: **Time-course trajectories of A549 treated with EGF.** Figure shows that our pipeline resulted in very variable and ambiguous trajectories for this case perhaps due to the lack of proper EMT induction via EGF on A549 cell line.

Preprint not peer reviewed



This preprint research paper has not been peer reviewed. Electronic copy available at: https://ssrn.com/abstract=4318055

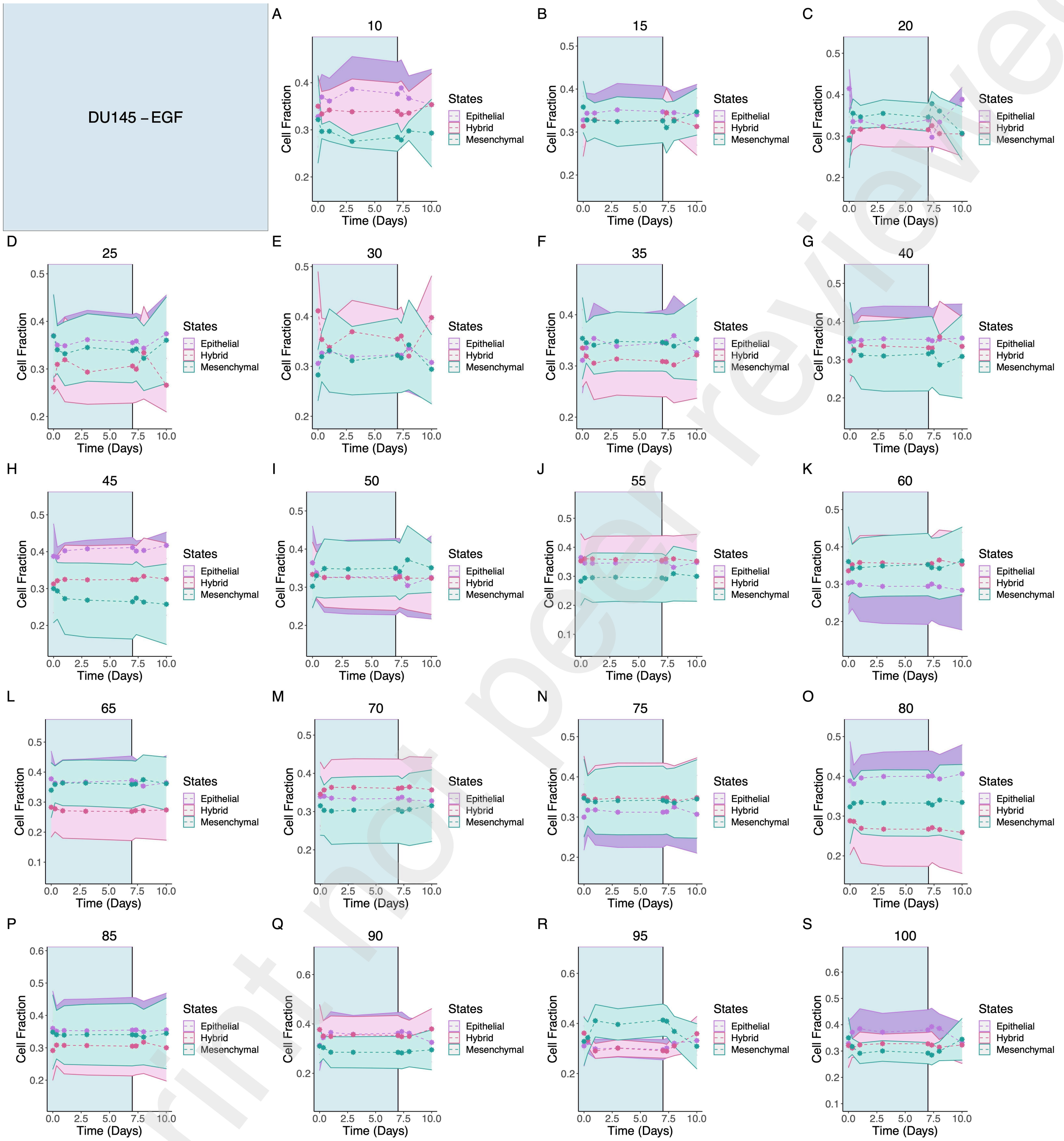


Figure SI.13: **Time-course trajectories of DU145 treated with EGF.** Figure shows that our pipeline resulted in very variable and ambiguous trajectories for this case perhaps due to the lack of proper EMT induction via EGF on DU145 cell line.



Preprint not peer reviewed

This preprint research paper has not been peer reviewed. Electronic copy available at: https://ssrn.com/abstract=4318055

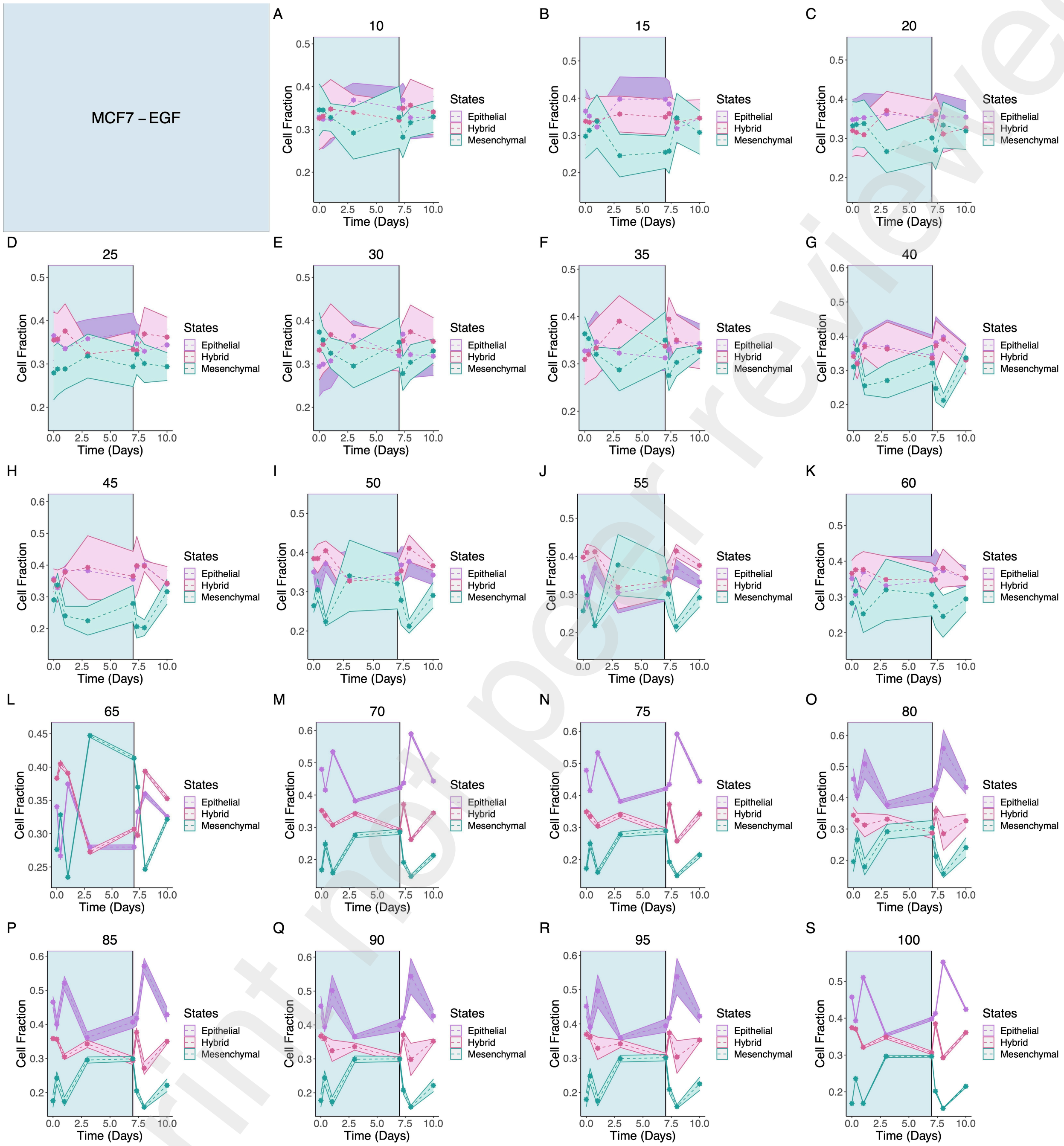


Figure SI.14: **Time-course trajectories of MCF7 treated with EGF.** Figure shows that our pipeline resulted in very variable and ambiguous trajectories for this case perhaps due to the lack of proper EMT induction via EGF on MCF7 cell line.

Preprint not peer reviewed



This preprint research paper has not been peer reviewed. Electronic copy available at: https://ssrn.com/abstract=4318055

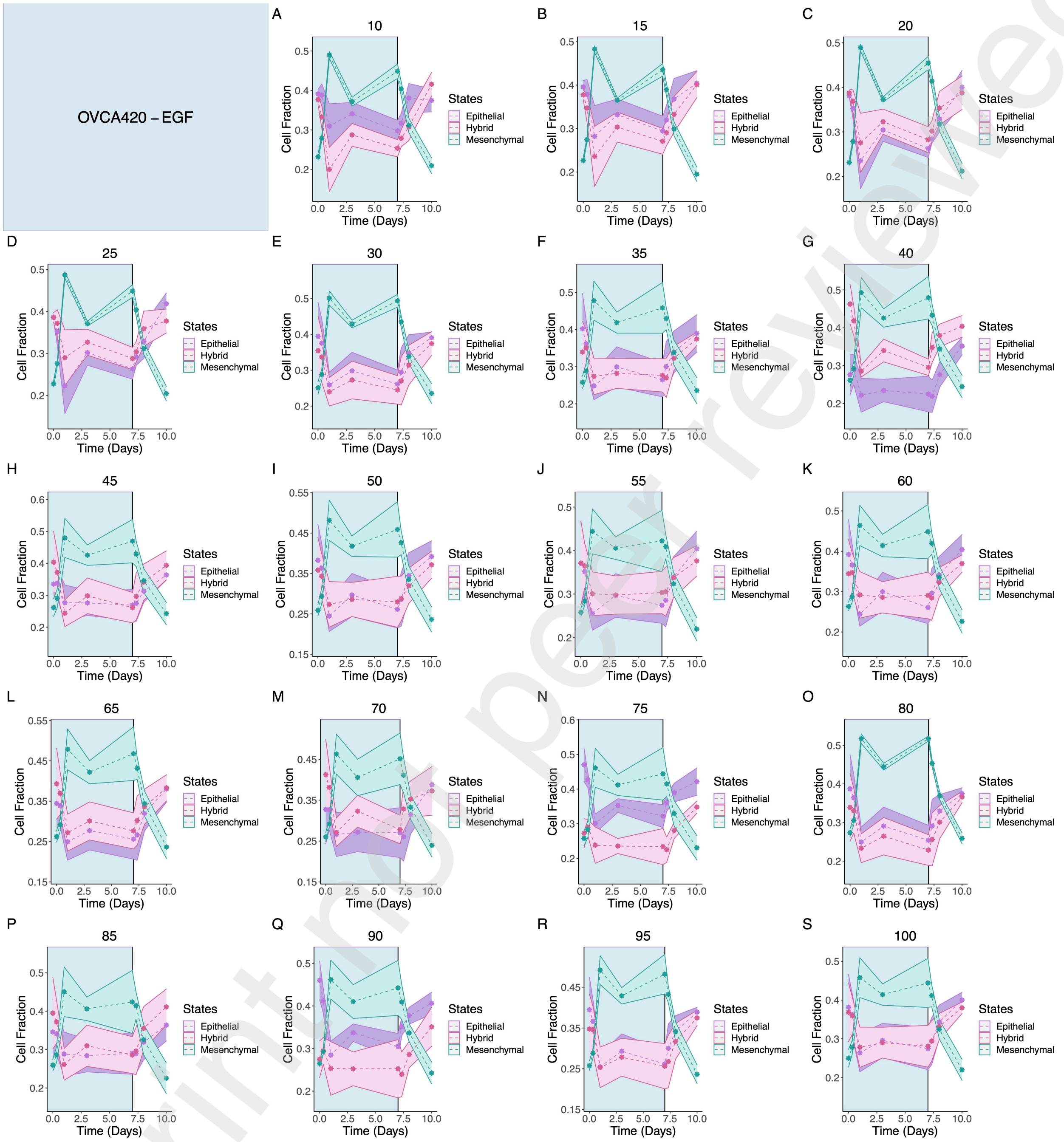


Figure SI.15: **Time-course trajectories of OVCA420 treated with EGF.** Figure shows that our pipeline overall predicted a rise in the mesenchymal state following treatment with EGF which falls and gives rise to hybrid and mesenchymal trajectories following treatment withdrawal.

Preprint not peer reviewed



This preprint research paper has not been peer reviewed. Electronic copy available at: https://ssrn.com/abstract=4318055

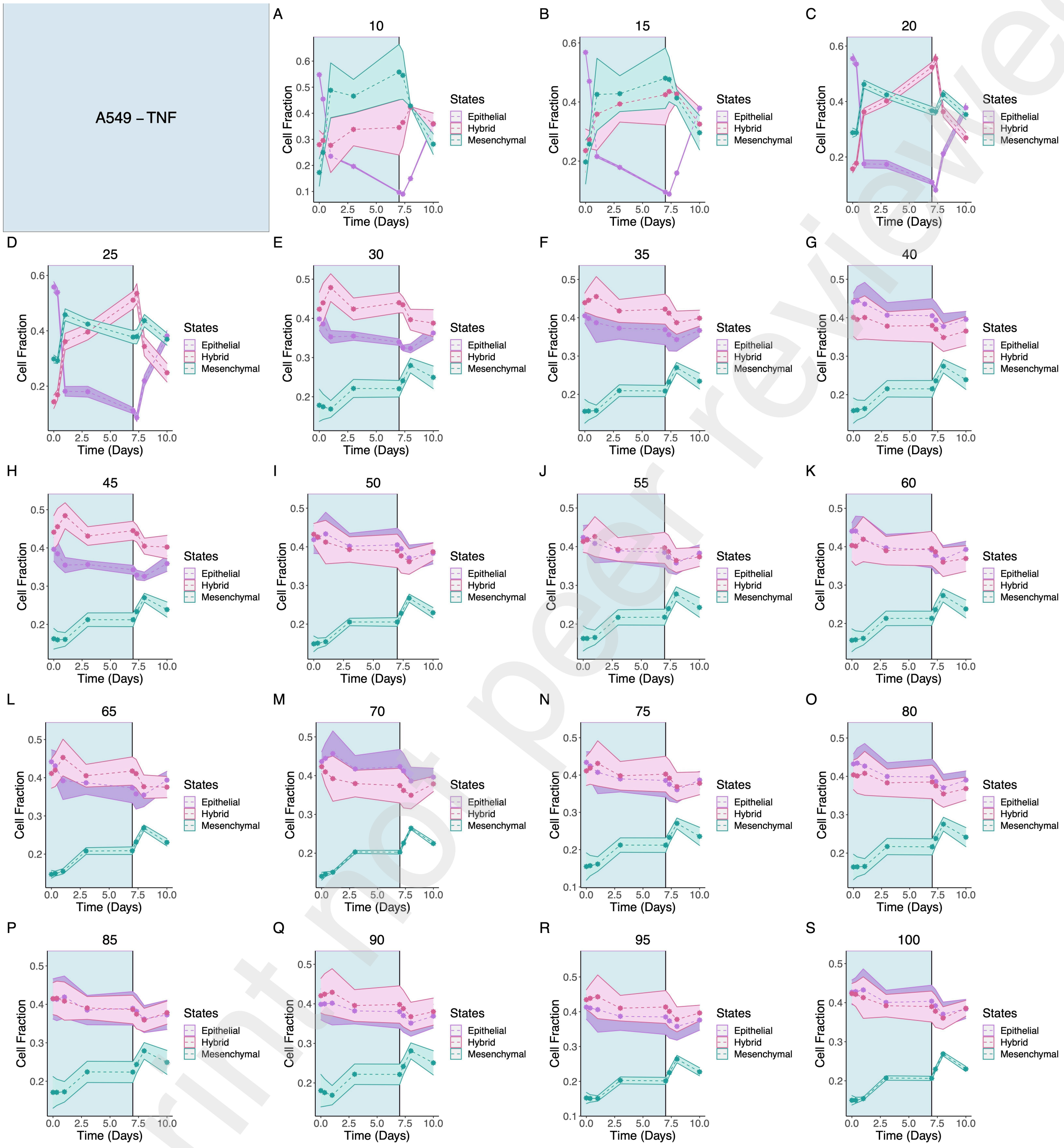


Figure SI.16: **Time-course trajectories of A549 treated with TNF.** Figure shows that our pipeline predicted a rise in the mesenchymal state following treatment with TNF which falls and gives rise to hybrid and mesenchymal trajectories following treatment withdrawal.



Preprint not peer reviewed

This preprint research paper has not been peer reviewed. Electronic copy available at: https://ssrn.com/abstract=4318055

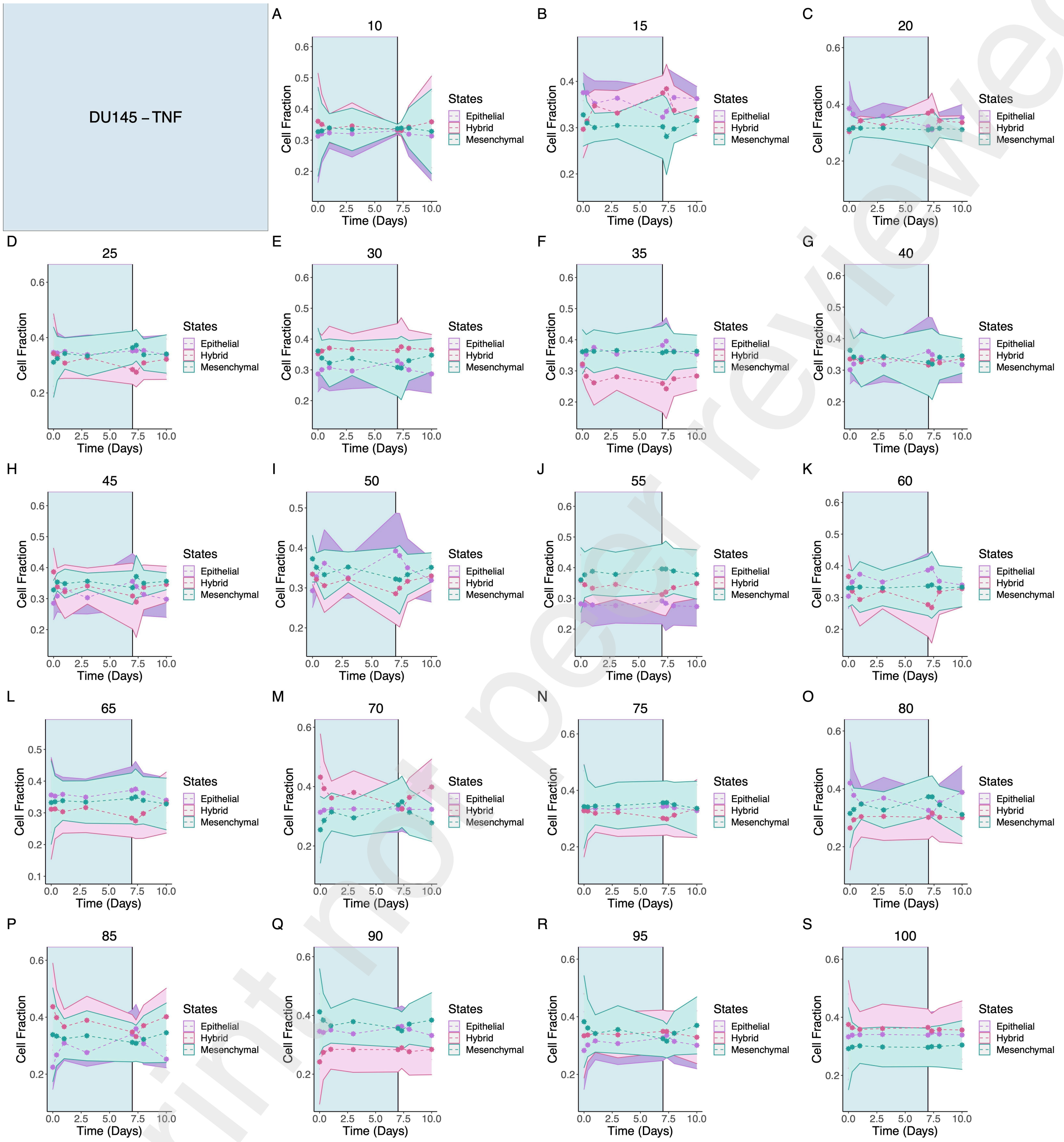


Figure SI.17: **Time-course trajectories of DU145 treated with TNF.** Figure shows that our pipeline overall predicted a rise in the mesenchymal state following treatment with TNF which falls and gives rise to hybrid and mesenchymal trajectories following treatment withdrawal for the lower cutoffs.

Preprint not peer reviewed



This preprint research paper has not been peer reviewed. Electronic copy available at: https://ssrn.com/abstract=4318055

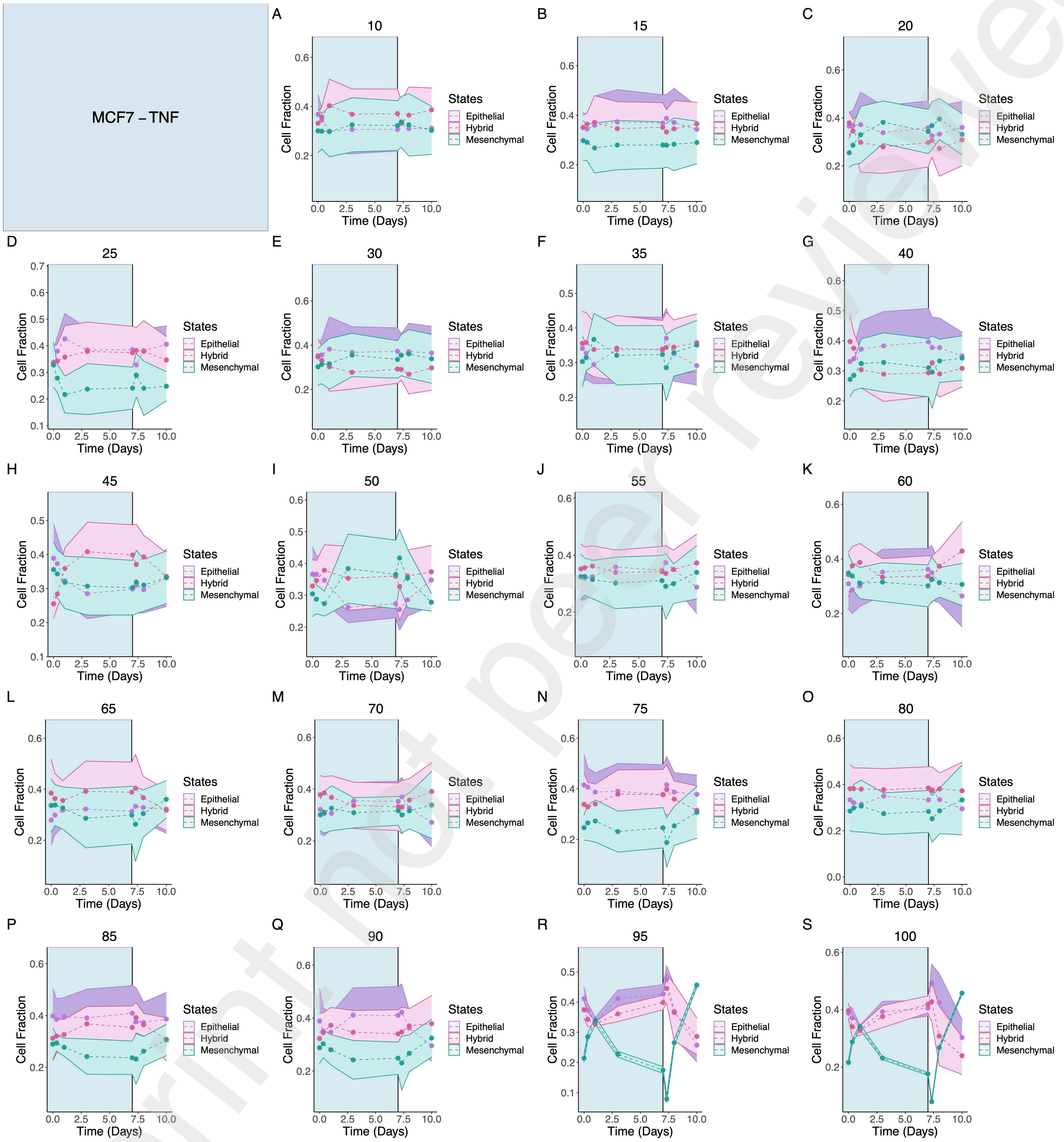


Figure SI.18: **Time-course trajectories of MCF7 treated with TNF.** Figure shows that our pipeline failed to infer transitions between trajectories reminiscent of EMT and MET.



Preprint not peer reviewed

This preprint research paper has not been peer reviewed. Electronic copy available at: https://ssrn.com/abstract=4318055

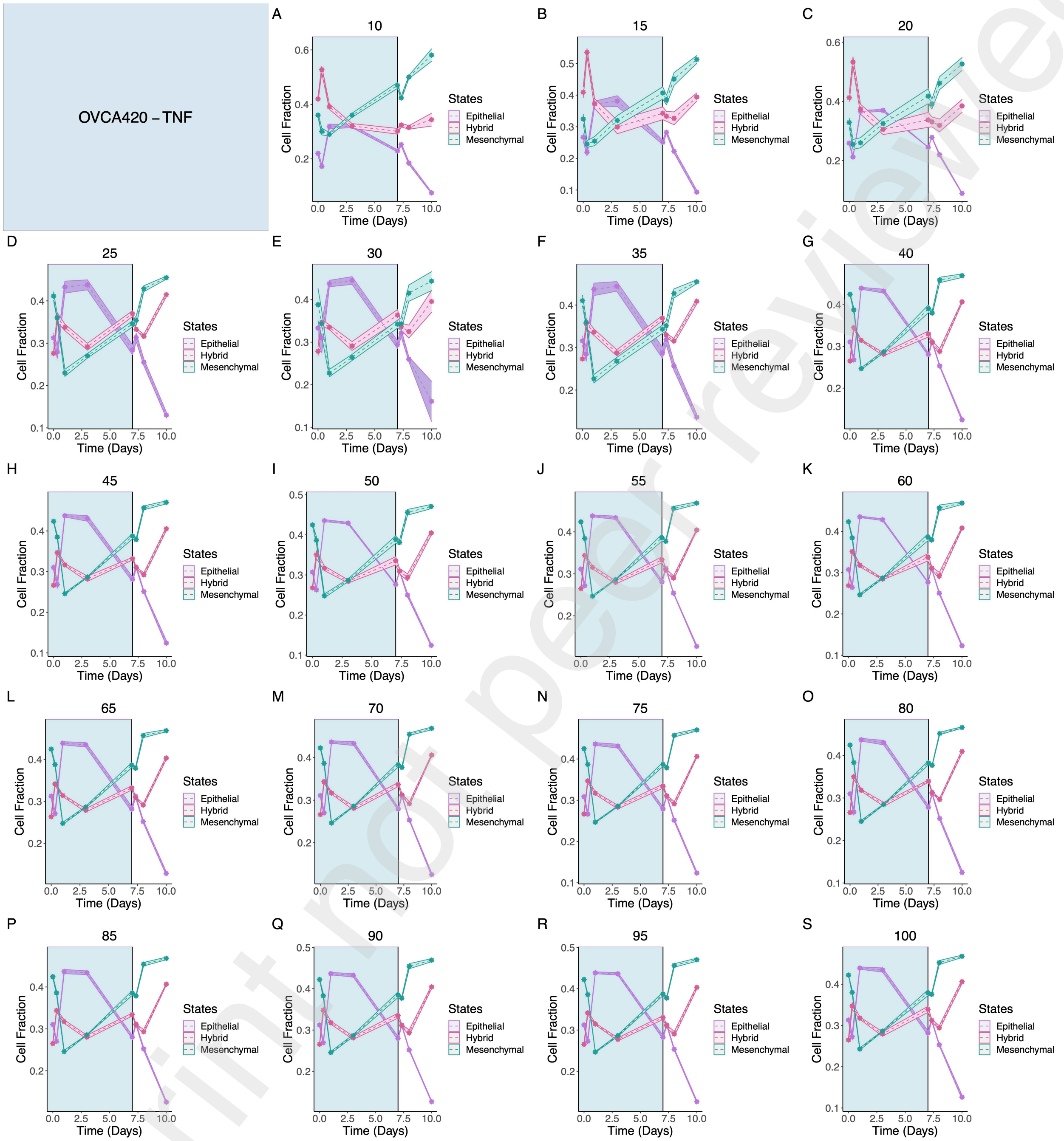


Figure SI.19: **Time-course trajectories of OVCA420 treated with TNF.** Figure shows that our pipeline may suggest an asynchronous EMT which continues after treatment withdrawal.



Preprint not peer reviewed

This preprint research paper has not been peer reviewed. Electronic copy available at: https://ssrn.com/abstract=4318055

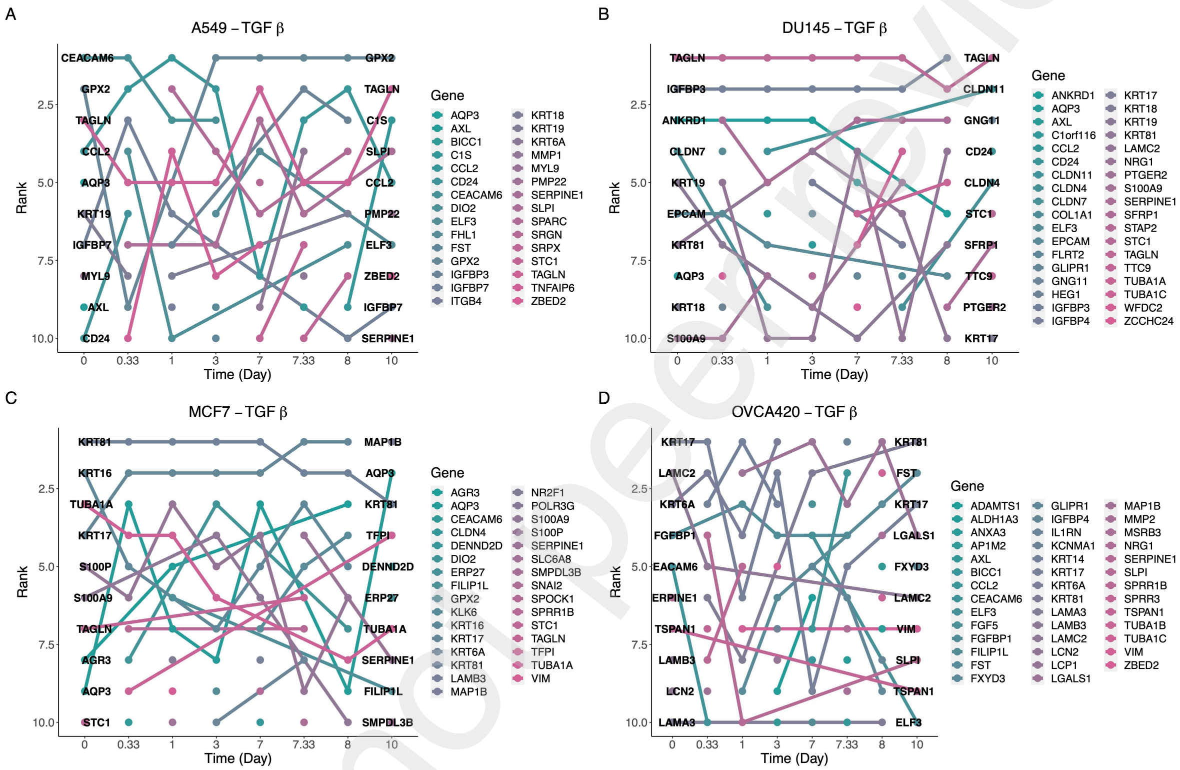


Figure SI.20: Plot showing how the top 10 variable genes switch rankings over time for TGF$\beta$ cases. Among this cohort, the OVCA420 cases has the most number of genes switching places.

.

Preprint not peer reviewed



This preprint research paper has not been peer reviewed. Electronic copy available at: https://ssrn.com/abstract=4318055

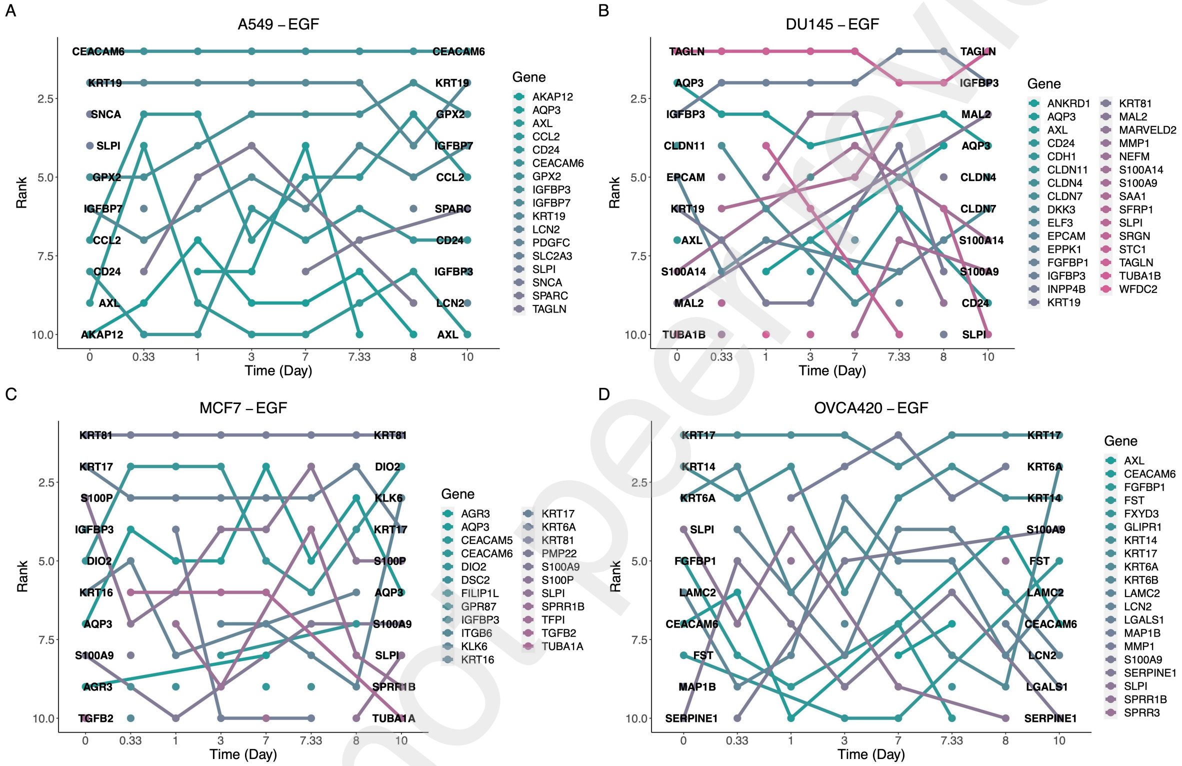


Figure SI.21: Plot showing how the top 10 variable genes switch rankings over time for EGF cases. It is evident that less genes switch places over time compared to the TGF$\beta$ cohort.

.



Preprint not peer reviewed

This preprint research paper has not been peer reviewed. Electronic copy available at: https://ssrn.com/abstract=4318055

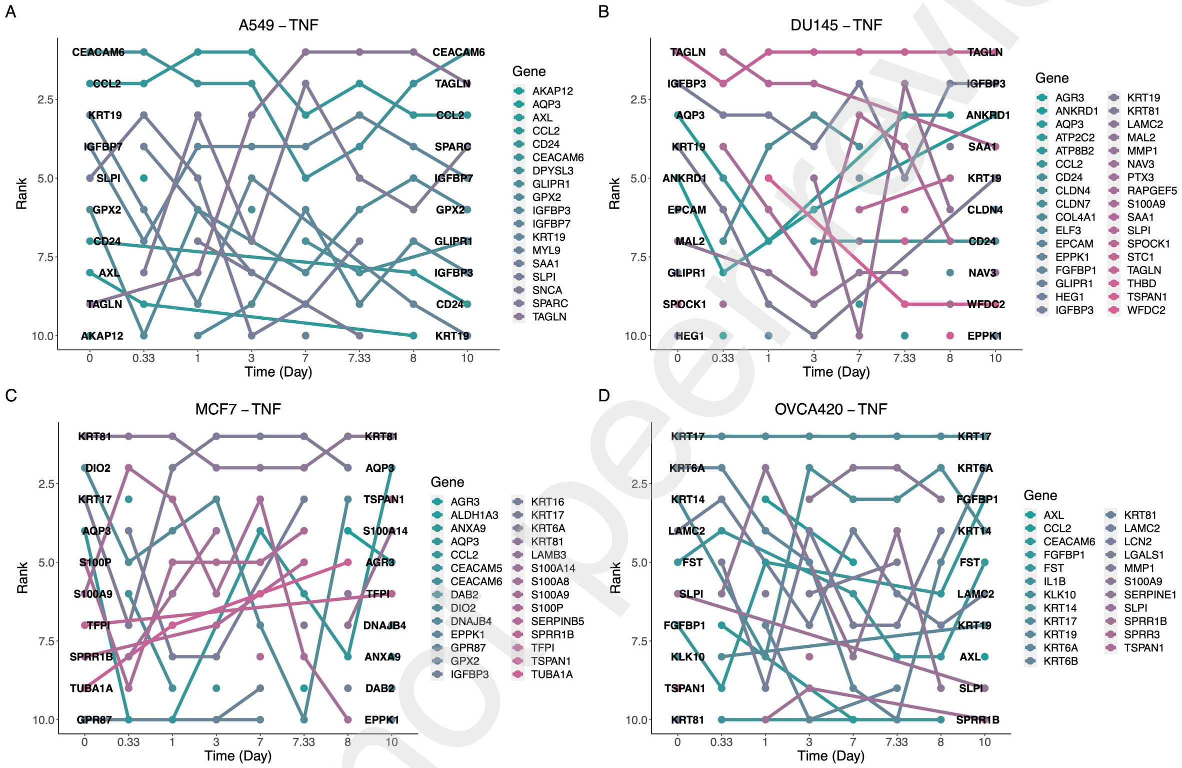


Figure SI.22: Plot showing how the top 10 variable genes switch rankings over time for TNF cases. The number of genes switching places in rankings are similar to the EGF case.

.

Preprint not peer reviewed



This preprint research paper has not been peer reviewed. Electronic copy available at: https://ssrn.com/abstract=4318055

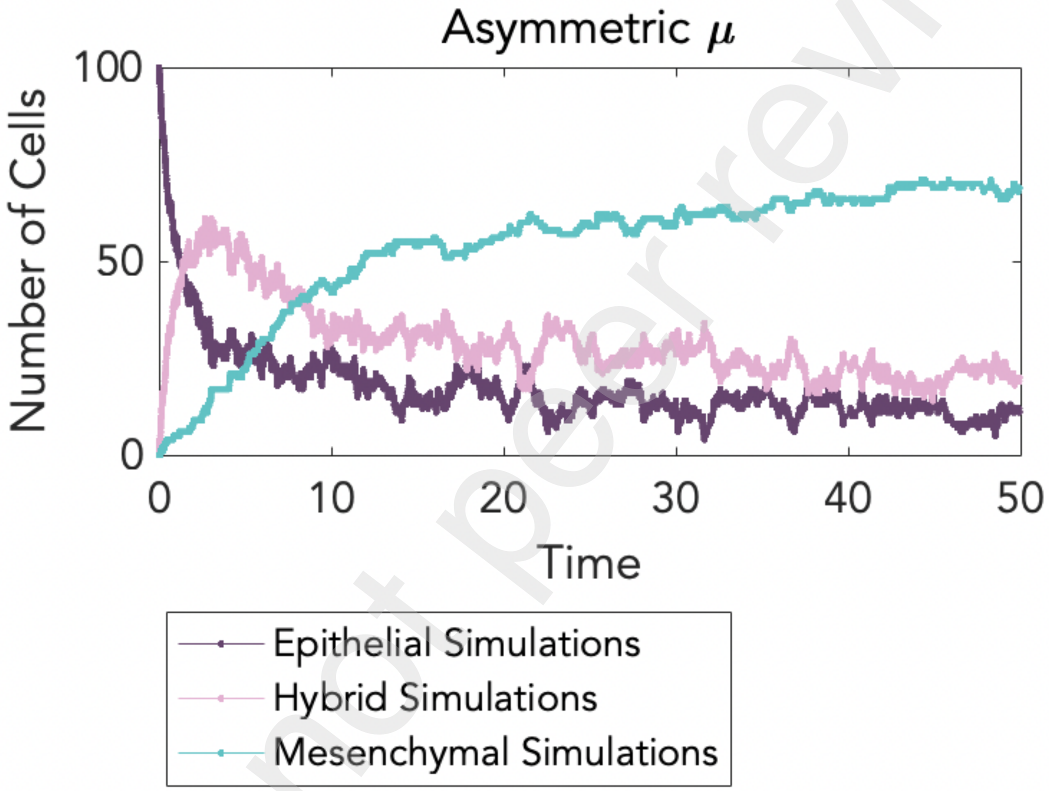


Figure SI.23: Plot depicts how simulations with asymmetric rates of transitions into H can recapitulate the dynamics of EMT with only one regime.



Preprint not peer reviewed

This preprint research paper has not been peer reviewed. Electronic copy available at: https://ssrn.com/abstract=4318055

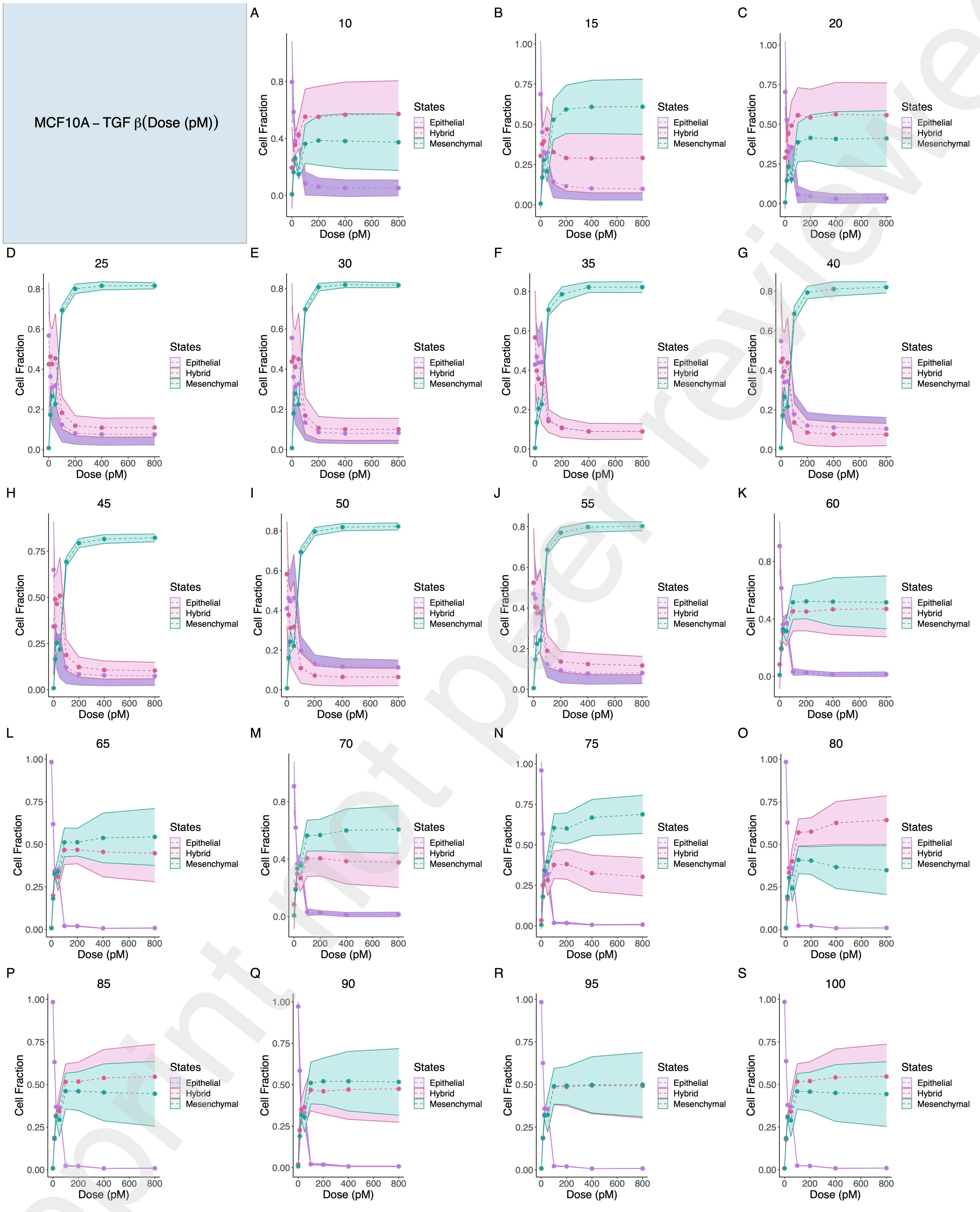

Figure SI.24: Plot shows the inferred trajectories for different cutoffs of highly variable genes for the steady state of MCF10A cell line treated with various doses of TGF$\beta$.



Preprint not peer reviewed

This preprint research paper has not been peer reviewed. Electronic copy available at: https://ssrn.com/abstract=4318055

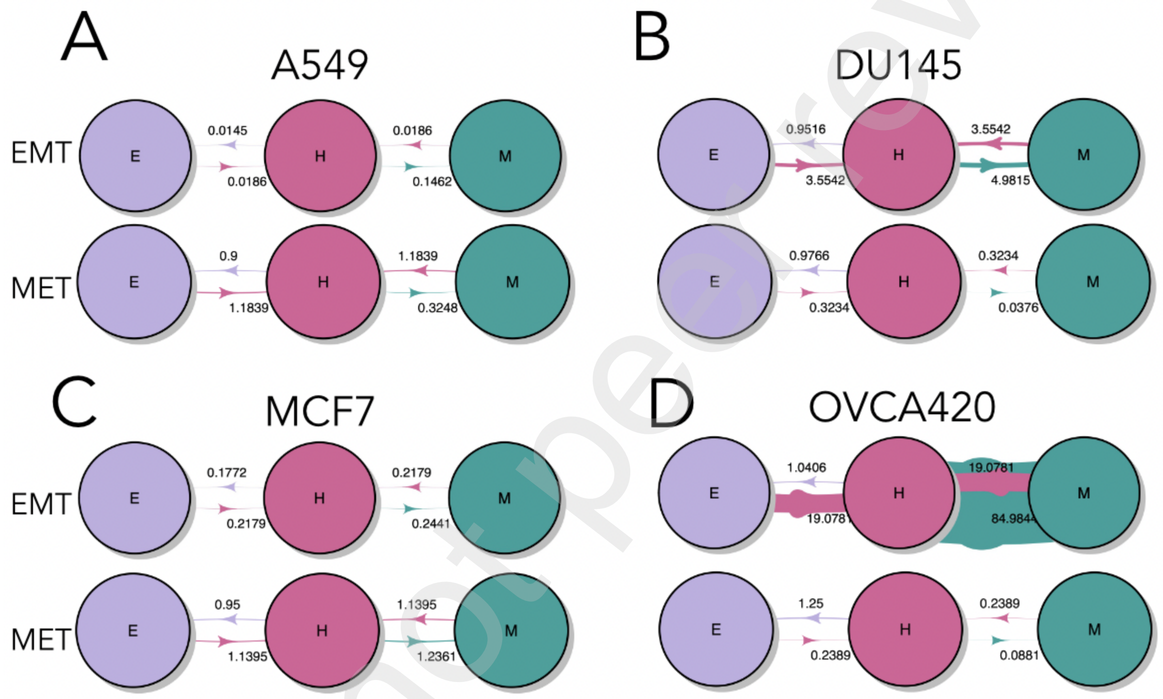


Figure SI.25: Figure shows the transition rates from and into states during EMT and MET obtained from fitting Markov chains to time-dependent data.

Preprint not peer reviewed



This preprint research paper has not been peer reviewed. Electronic copy available at: https://ssrn.com/abstract=4318055


Preprint not peer reviewed



This preprint research paper has not been peer reviewed. Electronic copy available at: https://ssrn.com/abstract=4318055
